# A Novel and Evolutionarily Conserved Metabolic Axis for Efficient Hepatic Lipid Catabolism Governed by Mfn2-Hsl-Mediated Mitochondria-Lipid Droplet Coupling

**DOI:** 10.64898/2026.09.28.754920

**Authors:** Ying-Jia Chi, Ze-Xuan Liu, Nan-Yun Hu, Guang-Li Feng, Yu-Feng Song

## Abstract

Inter-organelle contact between mitochondria and lipid droplets (LDs) is crucial for hepatic lipid homeostasis, but the tethering complex underlying this interaction remains unclear. To address this, we aimed to characterize the molecular machinery governing mitochondria-LD coupling and evaluate the evolutionary conservation of its metabolic functions. Using hepatocyte models from multiple species and *in vivo* experiments in yellow catfish with differential lipid and creatine diets, we explored the mechanism and evolutionary conservation of mitochondria-LD coupling. The key findings were: (1) A novel complex formed by mitofusin 2 on mitochondria and hormone-sensitive lipase on lipid droplets *via* specific residues orchestrates efficient hepatic lipid catabolism by mediating mitochondria-LD coupling; (2) Creatine enhances this coupling by transcriptionally upregulating *mfn2 via hnf4α*, thereby promoting peridroplet mitochondria formation and alleviating hepatic lipid accumulation; (3) The Mfn2-phosphorylated HSL axis is evolutionarily conserved across fish and mammals, representing a potential therapeutic target for hepatic steatosis.

## 1. INTRODUCTION

Lipid droplets (LDs) and mitochondria are two ubiquitous and essential organelles in eukaryotic cells, whose dynamic interactions play a pivotal role in maintaining lipid homeostasis,^1^ making this interplay a cutting-edge research focus in lipid metabolic studies.^2^ The canonical lipolysis-oxidation cascade involves: (1) LDs acting as the primary storage depot for neutral lipids, releasing free fatty acids (FFAs) *via* lipase-mediated lipolysis; (2) FFAs being transported to the mitochondrial matrix; and (3) undergoing FFAs *β*-oxidation to ultimately generate ATP.^3^ Thus, theoretically, the efficiency of this metabolic flux likely depends on LDs-mitochondria spatial proximity.^4^ Their membrane contact sites could significantly reduce FFA translocation distance.^5^ Recent studies, including the work published in Nature,^6^ demonstrated that LD-associated mitochondria significantly enhance cellular fatty acid *β*-oxidation capacity. We therefore hypothesize that a specialized, structurally coupled LDs-mitochondria metabolic channel that enables rapid and efficient lipid catabolism, although systematic investigations into this potential mechanism remain scarce. Given the global epidemic of hepatic steatosis,^7^ elucidating the molecular basis of this proposed LDs-mitochondria coupling-mediated and efficient lipid catabolic channel may provide novel therapeutic targets for precision interventions against metabolic disorders.

Mitochondrial fusion facilitates the expansion of the mitochondrial network, a process theorized to augment metabolic capacity for fatty acid *β*-oxidation (FAO).^8^ Central to this adaptive response is mitofusin 2 (Mfn2),^9^ which exhibits a dual functional repertoire: it serves not only as a core mediator of mitochondrial fusion but also as a critical structural scaffold enabling lipid droplet-mitochondria coupling.^10^ This bifunctional nature positions Mfn2 as a putative master regulator capable of enhancing FAO efficiency through spatial and functional coordination between the two organelles.^10^ From a mechanistic perspective, the surface of LDs serves as a platform for the enrichment of lipolytic enzymes, with lipolysis constituting the rate-limiting step in supplying FFAs to mitochondrial FAO.^11^ Our prior work established that Mfn2 potentiates upstream LDs lipolytic metabolism *via* facilitating mitochondrial-LD coupling;^12^ nevertheless, the precise molecular underpinnings remain incompletely characterized. A key unresolved question concerns the identity and composition of the protein complex, potentially organized around Mfn2 and LD-resident protein interactions, that concurrently sustains the physical LD-mitochondria interface and exerts regulatory control over both upstream LD lipolysis and downstream mitochondrial *β*-oxidation. Elucidating this mechanism would help define a structured, coupling-enabled pathway for accelerated lipid catabolism. Moreover, should such a dedicated metabolic channel exist, it becomes compelling to investigate which nutritional factors are capable of activating this pathway to mitigate hepatic steatosis. Resolving these aspects is expected to provide a conceptual advance toward nutrition-based strategies for the management of hepatic steatosis.

Creatine, a well-characterized nutritional supplement, has been widely recognized for its ability to enhance mitochondrial fatty acid oxidation,^13^ holding significant value in sports medicine and nutrition science.^14^ Emerging evidence suggests that creatine may also play a role in mitigating hepatic steatosis and restoring lipid homeostasis;^15^ however, the underlying molecular mechanisms remain unclear. Given coupling is a potential driver of mitochondrial FA oxidation,^16^ we are interested in an open issue: is there a novel mechanism of creatine elevating intracellular FAO capacity by strengthening mitochondria-lipid droplet coupling, thereby attenuating the progression of hepatic steatosis?

Mitochondria-LD coupling represents a structurally and functionally conserved inter-organelle communication system that has been extensively documented across diverse vertebrate species.^17,18^ Mfn2 also exhibits remarkable evolutionary conservation in both sequence and function among vertebrates.^19,20^ However, whether the fundamental mechanism of Mfn2-mediated mitochondria-LD coupling and its regulatory influence on lipolysis-coupled fatty acid oxidation maintain evolutionary conservation remains an important unresolved question in organelle biology and metabolic evolution. To address this knowledge gap, we implemented a phylogenetically stratified research strategy, progressing systematically from basal to derived vertebrate lineages, based on the established principle that ancestral species often retain conserved regulatory modules.^21^ Our investigation commenced with the teleost fish *Pelteobagrus fulvidraco*, a phylogenetically basal vertebrate that has emerged as a robust model for lipid metabolism research.^12,22,23^ In this system, we first identified hormone-sensitive lipase (HSL) as a crucial LD-resident protein and key lipolytic enzyme^24^ that physically interacts with Mfn2, and systematically deciphered how the Mfn2-HSL axis coordinates mitochondria-LD coupling to potentiate lipolysis and subsequent energy mobilization. Significantly, we precisely mapped the specific amino acid residues within both Mfn2 and HSL that govern mitochondria-LD tethering strength, thereby revealing strategic targets for therapeutic intervention at this metabolic interface. The evolutionary conservation of this mechanism was subsequently verified in higher vertebrate models, establishing its broad phylogenetic relevance. Furthermore, we investigated creatine supplementation could rescue hepatic lipid metabolic dysregulation through Mfn2/HSL-dependent mitochondria-LD communication, thereby proposing a novel targeted nutritional intervention strategy for hepatic steatosis.

## 2. RESULTS

### 2.1. Creatine ameliorates lipid overload-induced hepatic steatosis by promoting both hepatic triglyceride hydrolysis and fatty acid *β*-oxidation

In the present study, all groups maintained a 100% survival rate. Detailed growth performance, morphological indices, and feed utilization are shown in Table S1 (Supporting Information). Compared with the control group, lipid overload significantly decreased WG and SGR. Notably, dietary creatine supplementation resulted in significant elevations in WG and SGR, coupled with a reduced FCR, compared to other groups. These results suggest that dietary creatine supplementation improves growth performance and feed utilization in yellow catfish.

Results from Oil Red O and H&E staining demonstrated that in the lipid overload group, hepatic triglyceride deposition occupied over 25% of the tissue area, successfully establishing hepatic steatosis, while dietary creatine supplementation significantly alleviated lipid overload-induced hepatic steatosis, evidenced by the fact that compared with the lipid overload group, the triglyceride staining area in the lipid overload plus creatine group decreased by approximately 6.05% (Figure 1A-H). Generally, hepatic lipid accumulation reflects the balance among dietary lipid absorption, de novo lipogenesis, and lipid catabolism *via* lipolysis and *β*-oxidation. Interestingly, further qPCR analysis demonstrated that enhanced hepatic lipolysis and subsequent *β*-oxidation are likely the primary mechanisms through which creatine alleviates hepatic steatosis, evidenced by the up-regulated expression of genes involved in lipolysis and *β*-oxidation (*g6pd, 6pgd, acca, fas, dgat1, srebp1, atgl, hsl, mgl, acsl1, cpt1, cpt2, lcad, mcad, scad* and *echs*) in lipid overload + creatine group (Figure 1I). Further support is provided by the activity of key enzymes in lipolysis and *β*-oxidation and their metabolic intermediates (Figure 1J-R and S1A, B). Collectively, these results suggest that hepatic lipolysis and subsequent *β*-oxidation are likely the principal mechanisms driving creatine-mediated alleviation of hepatic steatosis.

**Fig. 1.**
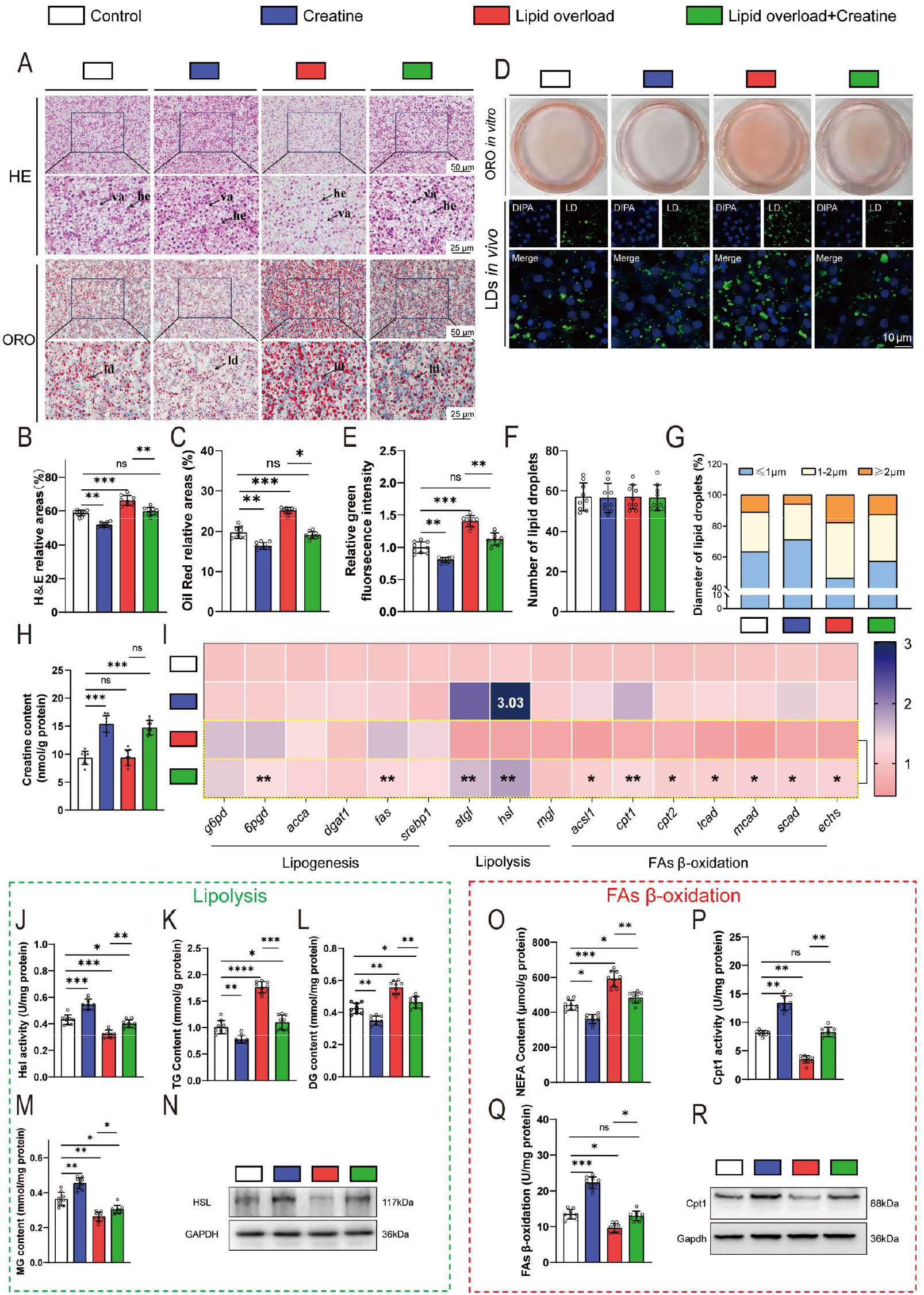
Creatine ameliorates lipid overload-induced hepatic steatosis by promoting both hepatic triglyceride hydrolysis and fatty acid *β*-oxidation. A) Representative images of liver tissues stained by H&E and oil red O (*in vivo* experiments). 200 × magnification. scale bars, 50 μm; HE, hepatocytes. VA, vacuoles. B, C) Relative areas for hepatic vacuoles after H&E staining and LDs after oil red O staining, respectively. D) Representative images of hepatocyte stained by oil red O and Bodipy, respectively (*in vivo* and *in vitro* experiments). E) Relative green fluorescence intensity in panel D. F) Number of lipid droplets in panel D. G) Quantification of lipid droplet diameter in panel D. H) Creatine content (*in vivo* experiments). I) The mRNA levels of genes involved in hepatic lipid homeostasis (*in vivo* experiments). J) HSL activity (*in vivo* experiments). K) TG content (*in vivo* experiments). L) DG content (*in vivo* experiments). M) MG content (*in vivo* experiments). N) Western blot analysis of HSL (*in vivo* experiments). O) NEFA content (*in vivo* experiments). P) Cpt1 activity of isolated mitochondria from tissues (*in vivo* experiments). Q) FAs *β*-oxidation rate of isolated mitochondria from tissues (*in vivo* experiments). R) Western blot analysis of Cpt1 (*in vivo* experiments). Lipid overload: high-fat diet (HFD) for *in vivo* in A, B, C, E, F, G, H, I, J, K, L, M, N, O, P panels; FA (OA:PA/1:1) incubate for *in vitro* in D panels. For western blots, n=3 three biological replicates of one treatment. For the other data, *in vivo*, n = 9, 3 tanks for per diet with 3 fish randomly from each tank; *in vitro*, n = 9, including three biological replicates and three technical replicates per biological replicate. Statistical significance was defined as a p-value < 0.05. Significance levels were denoted as follows: * P<0.05, ** P<0.01, *** P<0.001, and **** P<0.0001. Non-significant differences were labeled "ns".

### 2.2. Dietary creatine promotes mitochondrial fusion and then increases PDM formation by activating the *hnf4α*-binding element in the mfn2 promoter

Since dietary creatine enhances fatty acid *β*-oxidation, and also given a mitochondrial process dependent on mitochondrial quantity and dynamics, we investigated whether creatine promotes *β*-oxidation by modulating mitochondrial quantity or dynamics. Flow cytometry, mtDNA copy number, and Tom20 protein expression analyses revealed that neither dietary creatine nor lipid overload significantly affected mitochondrial quantity (Figure 2A, B and S2A-C). Confocal microscopy demonstrated that lipid overload increased mitochondrial fragmentation, whereas creatine treatment elevated the proportion of elongated mitochondria (Figure 2C, D), suggesting that dietary creatine activates mitochondrial fusion without altering mitochondrial quantity. Further investigation indicated that, among key mitochondrial dynamics-related proteins, Mfn2 plays a critical role in creatine-activated mitochondrial fusion (Figure 2E and S2D, E). To elucidate the molecular mechanism underlying the transcriptional regulation of *mfn2*, bioinformatic predictions of putative transcription factors binding to the *mfn2* promoter region were performed using three public databases: Harmonizome, JASPAR, and hTFtarget (Figure 2F). The candidate transcription factors were ranked according to their predicted binding scores in descending order (Figure 2G-I). Intersection analysis of the three datasets identified *tfap2c*, *hnf4α*, *foxo3*, and *gata2* as the transcription factors with the highest binding potential to the *mfn2* promoter. Based on these in silico analyses, qPCR was subsequently conducted for experimental validation. The results demonstrated that the transcriptional level of *hnf4α* exhibited the most significant change following creatine treatment (Figure 2J). Dual-luciferase reporter and electrophoretic mobility shift assays (EMSA) confirmed that creatine upregulates *mfn2* expression primarily through activation of the *hnf4α* cis-element within its promoter region (Figure 2K, L).

**Fig. 2.**
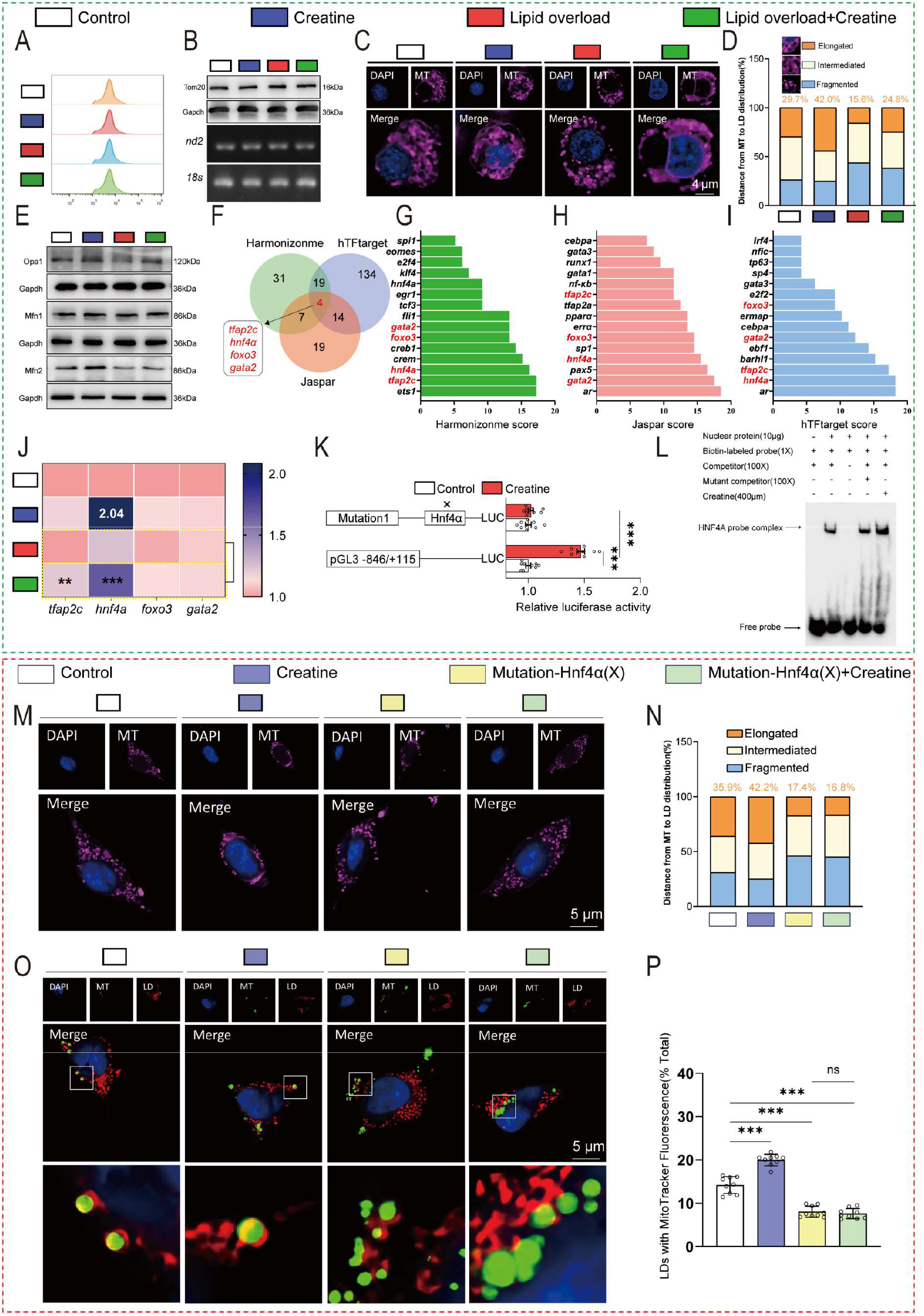
Dietary creatine promotes mitochondrial fusion and then increases PDM formation by activating the HNF4α-binding element in the mfn2 promoter. A) Flow cytometry analysis of mitochondrial content (*in vitro* experiments). B). Western blot analysis of Tom20 and mtDNA fold analysis (*in vivo* experiments). C) Confocal images of mitochondria (*in vitro* experiments). scale bars, 5 µm. D) Quantification of mitochondrial states (elongated: ≥1 μm, intermediate: 0.5-1 μm, and fragmented: ≤0.5 μm) in panel C. E) Western blot analysis of Opa1, Mfn1 and Mfn2 (*in vivo* experiments). F) Bioinformatics analysis identifies potential transcription factors driving *mfn2* expression. G) Prediction of potential transcription factors governing *mfn2* gene expression using the Harmonizonme. H) Prediction of potential transcription factors governing *mfn2* gene expression using the Jaspar. I) Prediction of potential transcription factors governing *mfn2* gene expression using the hTFtarget. J) The mRNA levels of genes (*in vivo* experiments). K) Sitemutation assay of *hnf4α* binding sites on mfn2-846 vector in 293T cells (*in vitro* experiments). L) EMSA of *hnf4*α binding sequence (*in vitro* experiments). M) Confocal images of mitochondria (*in vitro* experiments). scale bars, 5 µm. N) Quantification of mitochondrial states (elongated: ≥1 μm, intermediate: 0.5-1 μm, and fragmented: ≤0.5 μm) in panel M. O) Representative confocal microscopy image after MitoTracker Deep Red and BODIPY493/503 staining (*in vitro* experiments). P) The fusion rate between MitoTracker and LDs in panel O. Lipid overload: high-fat diet (HFD) for *in vivo* in B, E, J panels; FA (OA:PA/1:1) incubate for *in vitro* in A, C, D, K, L, M, N, O, P panels. For western blots, n=3 three biological replicates of one treatment. For the other data, *in vivo*, n = 9, 3 tanks for per diet with 3 fish randomly from each tank; *in vitro*, n = 9, including three biological replicates and three technical replicates per biological replicate. Statistical significance was defined as a p-value < 0.05. Significance levels were denoted as follows: * P<0.05, ** P<0.01, *** P<0.001, and **** P<0.0001. Non-significant differences were labeled "ns".

To rule out potential effects elicited by other biological functions of creatine, we constructed a hepatocyte model with site-directed mutation of the *hnf4α* binding element on the endogenous *mfn2* promoter, in which the core binding motif was mutated to block *hnf4α* recognition and binding. The results showed that after mutating this *hnf4α* binding site, creatine treatment no longer upregulated Mfn2 transcriptional and protein expression (Figure S2F, G), and the promotive effects of creatine on mitochondrial fusion and PDM formation were completely abolished (Figure 2M-P). Meanwhile, creatine-induced enhancements in lipid droplet lipolysis and mitochondrial fatty acid *β*-oxidation were also fully blocked in this mutant model (Figure S2H-L).

### 2.3 Dietary creatine promotes PDM formation with high fatty acid oxidation capacity

After the interaction between lipid droplets and mitochondria, the physical distance between the two organelles is narrowed. This physical alteration is considered a critical structural basis for material exchange. Accordingly, transmission electron microscopy (TEM) observations confirmed the colocalization of lipid droplets and mitochondria. Results revealed that creatine-enhanced, Mfn2-mediated mitochondrial fusion (Figure 3A, B) was accompanied by an increased overlap ratio between mitochondria and lipid droplet perimeters (Figure 3A-E). These observations were further corroborated by 2D/3D confocal imaging (Figure 3F-H).

**Fig. 3.**
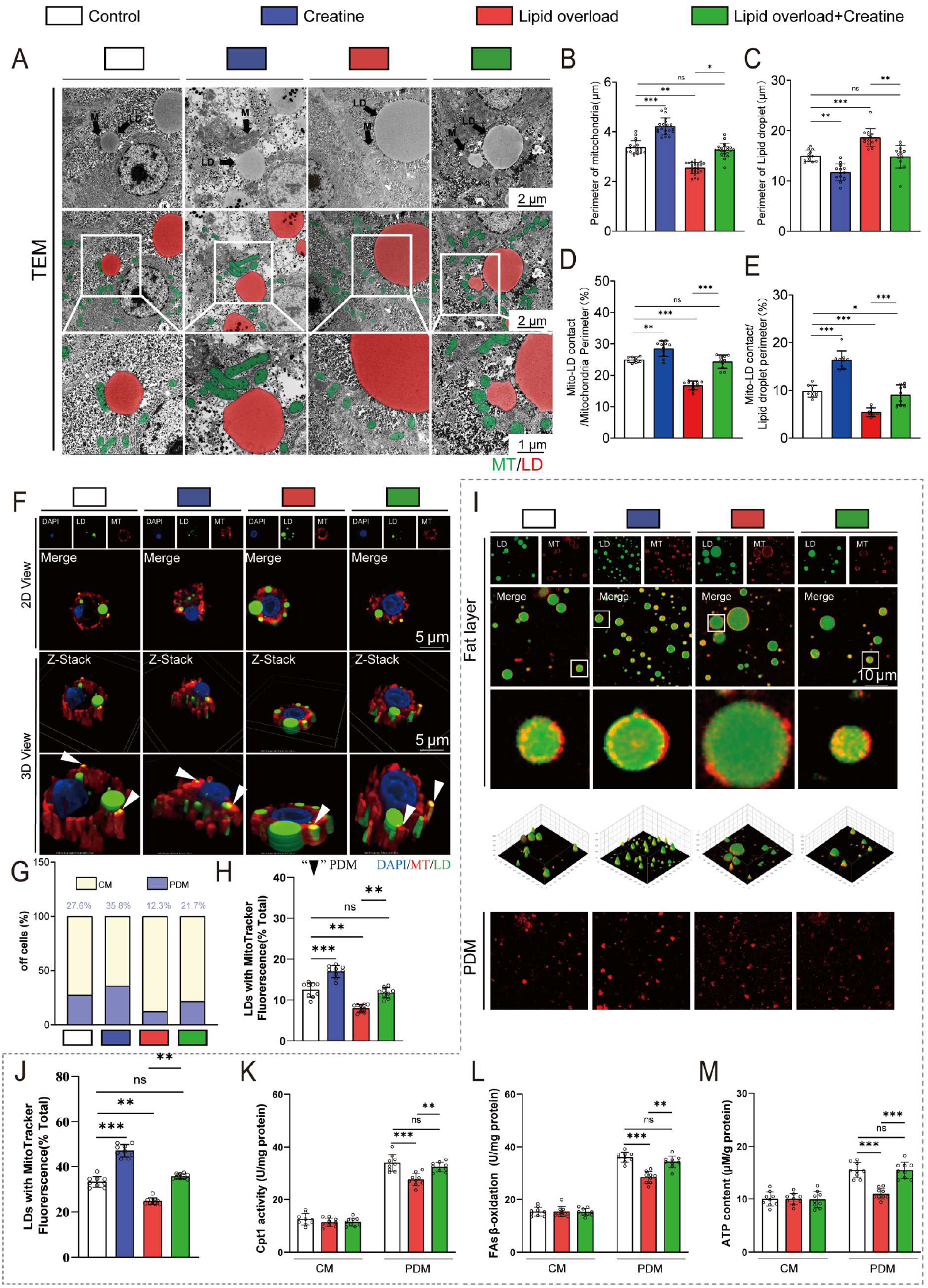
Dietary creatine promotes PDM formation with high fatty acid oxidation capacity. A) Representative images of hepatic ultrastructure (TEM, *in vivo* experiments). B) Perimeter of mitochondria in panel A. C) Perimeter of lipid droplet in panel A. D) Ratio of LD-mito overlapping perimeter to total mitochondrial perimeter in panel A. E) Ratio of LD-mito overlapping perimeter to total lipid droplet perimeter in panel A. F) Representative confocal microscopy image after MitoTracker Deep Red and BODIPY493/503 staining (*in vitro* experiments). G) Quantification of the content ratio of PDM and CM in panel F. H) The fusion rate between MitoTracker and LDs in panel F. I) Confocal images of PDM which were stripped from LDs (*in vivo* experiments). scale bars, 10 µm. J) The fusion rate between MitoTracker and LDs in panel I. K) Cpt1 activity of isolated PDM and CM from tissues (*in vivo* experiments). L) FAs *β*-oxidation rate of isolated PDM and CM from tissues (*in vivo* experiments). M) ATP content (*in vivo* experiments). Lipid overload: high-fat diet (HFD) for *in vivo* in A, B, C, D, E, I, J, K, L, M panels; FA (OA:PA/1:1) incubate for *in vitro* in F, G, H panels. For western blots, n=3 three biological replicates of one treatment. For the other data, *in vivo*, n = 9, 3 tanks for per diet with 3 fish randomly from each tank; *in vitro*, n = 9, including three biological replicates and three technical replicates per biological replicate. Statistical significance was defined as a p-value < 0.05. Significance levels were denoted as follows: * P<0.05, ** P<0.01, *** P<0.001, and **** P<0.0001. Non-significant differences were labeled "ns".

To explore the functional implications, we isolated and purified distinct mitochondrial subpopulations-cytosolic mitochondria (CM) and perilipin-enriched droplet-associated mitochondria (PDM). Our results showed that lipid overload significantly reduced the proportion of PDM in hepatocytes, whereas creatine treatment markedly increased it (Figure 3I, J). Comparative analysis of fatty acid *β*-oxidation capacity between CM and PDM demonstrated that PDM exhibited a higher *β*-oxidation rate (Figure 3K-M). Given the previously demonstrated enhancement of fatty acid *β*-oxidation by creatine, we propose that this effect is likely mediated through an increase in the intracellular proportion of PDM.

### 2.4. Mfn2 modulates the formation of lipid droplet-mitochondria coupling *via* its protein interaction with HSL

Having established that Mfn2-mediated mitochondrial fusion promotes LD-mitochondria coupling, we sought to delineate the underlying mechanism. As mitochondrial quantity is often linked to inter-organelle contacts, we first assessed whether *mfn2* knockdown or creatine treatment altered mitochondrial number. However, neither intervention significantly altered intracellular mitochondrial quantity, as confirmed by mtDNA copy number, flow cytometry analysis, and Tom20 protein expression (Figure S3A-F). Importantly, confocal microscopy revealed that *mfn2* knockdown markedly reduced the formation of mitochondria-lipid droplet contact structures (Figure 4A-B). To further verify this finding, we performed *in vitro* reconstitution assays with isolated mitochondria and lipid droplets. The results showed that creatine significantly promoted the physical interaction between mitochondria and LDs, whereas *mfn2* knockdown largely abolished this effect (Figure 4C-E), further confirming the essential role of Mfn2 in mediating this inter-organelle interaction.

**Fig. 4.**
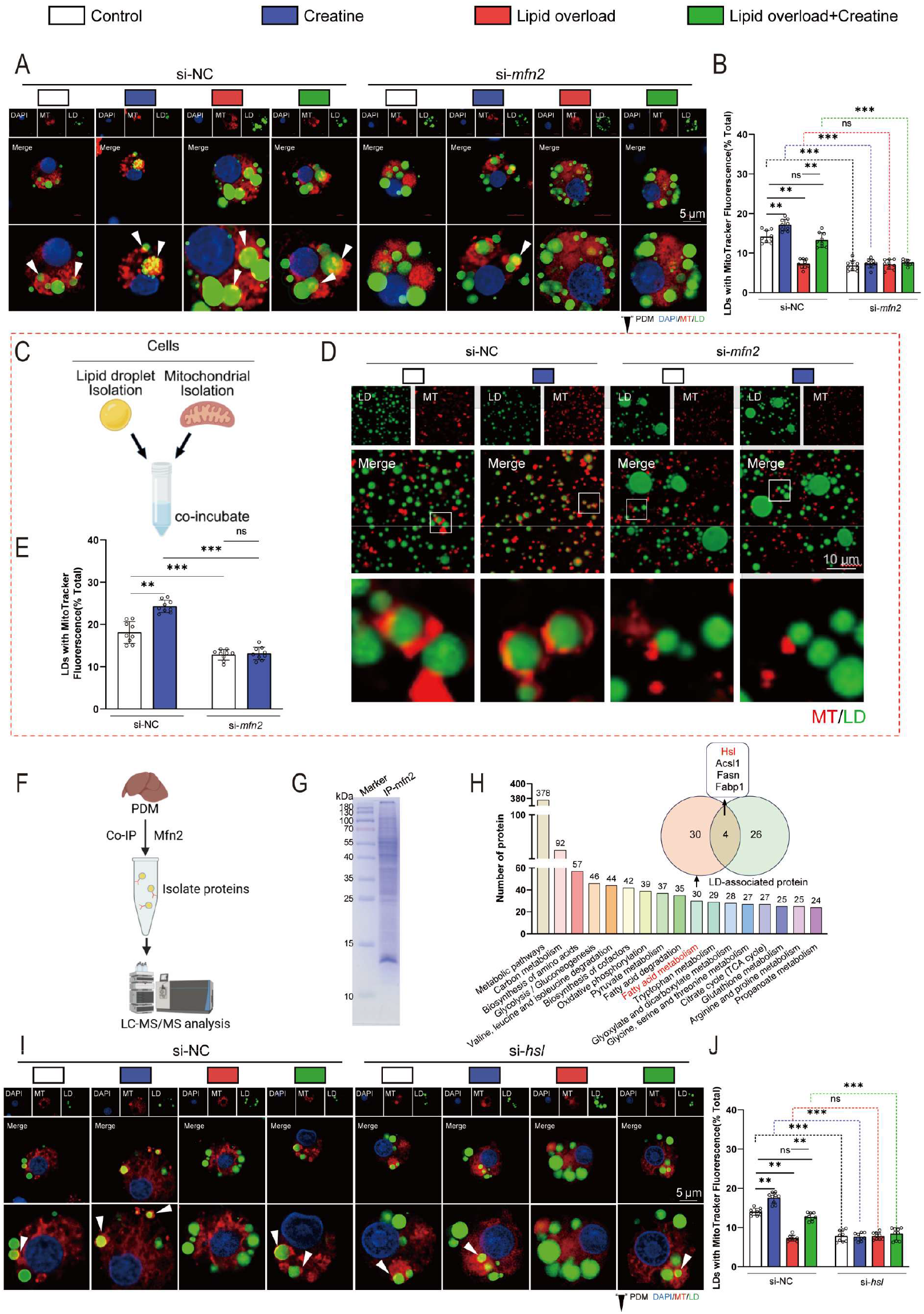
Mfn2 modulates the formation of lipid droplet-mitochondria coupling *via* its protein interaction with HSL. A) Representative confocal microscopy image after MitoTracker Deep Red and BODIPY493/503 staining (*in vitro* experiments). B) The fusion rate between MitoTracker and LDs in panel A. C) Schematic representation of the experimental design. D) Confocal images of PDM stripped from primary hepatocytes treated with si-NC or si-*mfn2* for 48 h scale bars, 10 µm (*in vitro* experiments). E) The fusion rate between MitoTracker and LDs in panel D. F) Schematic representation of the experimental protocols (*in vivo* experiments). G) Coomassie blue staining images for Mfn2-binding proteins (*in vivo* experiments). H) Kyoto Encyclopedia of Genes and Genomes (KEGG) analysis revealed the protein-enriched signaling pathways of the proteins interacting with Mfn2 (*in vivo* experiments). I) Representative confocal microscopy image after MitoTracker Deep Red and BODIPY493/503 staining (*in vitro* experiments). J) The fusion rate between MitoTracker and LDs in panel I. FA (OA:PA/1:1) incubate for *in vitro* in A, B, C, D, H, I panels. For all data, *in vivo*, n = 9, 3 tanks for per diet with 3 fish randomly from each tank; *in vitro*, n = 9, including three biological replicates and three technical replicates per biological replicate. Statistical significance was defined as a p-value < 0.05. Significance levels were denoted as follows: * P<0.05, ** P<0.01, *** P<0.001, and **** P<0.0001. Non-significant differences were labeled "ns".

Mitochondria-mediated organelle interactions typically require physical tethering between mitochondrial-resident proteins and proteins on partner organelles. To identify potential binding partners of Mfn2 on lipid droplets, we performed co-immunoprecipitation of Mfn2 from purified PDM fractions followed by LC-MS/MS analysis (Figure 4F, G). This approach identified 378 Mfn2-interacting proteins, including 4 candidates selected based on the intersection of Mfn2-binding proteins and LD proteins (Figure 4H). Notably, among these were HSL and other proteins critically involved in lipid metabolism. Since we had previously demonstrated that creatine promotes lipid droplet-mitochondria coupling along with the activation of mitochondrial *β*-oxidation and LD lipolytic, we focused on the role of Mfn2-HSL complex on lipid droplet-mitochondria coupling. Subsequent experiments showed that *hsl* knockdown, without affecting mitochondrial quantity, as evidenced by mtDNA copy number, Tom20 protein expression, and flow cytometry analysis (Figure S3G-L), similarly reduced lipid droplet-mitochondria coupling structures, and this was confirmed by confocal images (Figure 4I, J). These findings collectively demonstrate that Mfn2 modulates lipid droplet-mitochondria interactions through its specific protein-protein interaction with HSL.

### 2.5. Mfn2-HSL-mediated mitochondria-lipid droplet coupling establishes an efficient channel linking LD hydrolysis to mitochondrial fatty acid *β*-oxidation

Having established the necessity of Mfn2-HSL in maintaining mitochondria-LD coupling, we next investigated the functional impact of this inter-organelle association on both organelles. Following *hsl* knockdown, which disrupted LD-mitochondria tethering, we observed a significant alteration in mitochondrial dynamics characterized by increased mitochondrial fragmentation (Figure 5A, B). This structural disruption was accompanied by a marked reduction in mitochondrial fatty acid *β*-oxidation efficiency, as evidenced by decreased Cpt1 enzyme activity (Figure 5C, D) and supported by the observation that after *hsl* knockdown, C12-labeled fatty acids showed a marked reduction in their ability to transfer to mitochondria (Figure 5E, F). These findings indicate that HSL-mediated organelle coupling not only maintains physical linkage but also critically supports mitochondrial fusion dynamics and functional capacity for mitochondrial fatty acid *β*-oxidation.

**Fig. 5.**
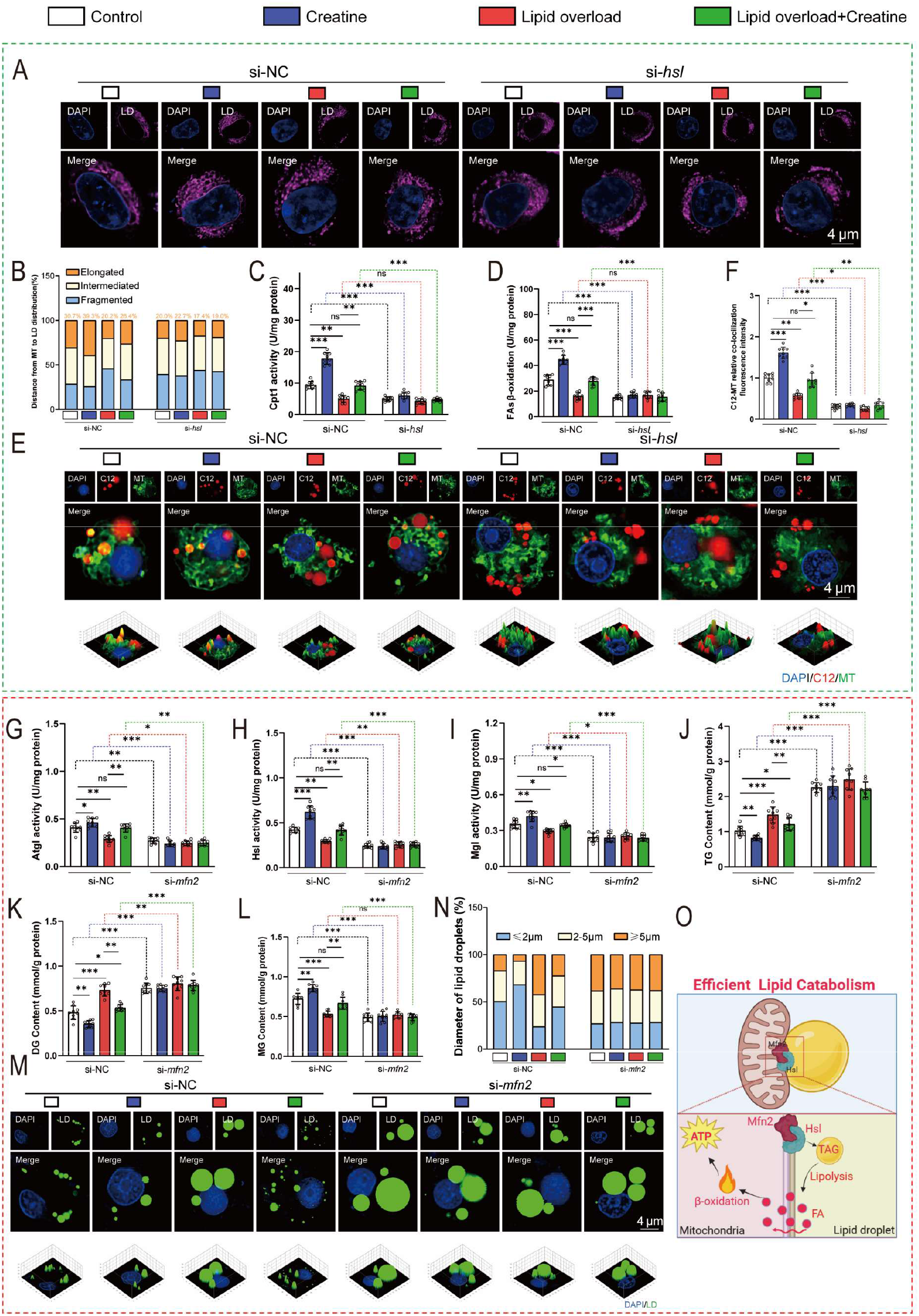
Mfn2-HSL-mediated mitochondria-lipid droplet coupling establishes an efficient channel linking LD hydrolysis to mitochondrial fatty acid *β*-oxidation (*in vitro* experiments). A) Confocal images of mitochondria. scale bars, 5 µm. B) Quantification of mitochondrial states (elongated: ≥1 μm, intermediate: 0.5-1 μm, and fragmented: ≤0.5 μm) in panel A. C) Cpt1 activity of isolated mitochondria from cells. D) FAs *β*-oxidation rate of isolated mitochondria from cells. E) Representative confocal images of mitochondria and Red C12 480/508, scale bars, 4µm. F) The fusion rate between mitochondria and Red C12 in panel E. G) Atgl activity. H) HSL activity. I) Mgl activity. J) TG content. K) DG content. L) MG content. M) Representative confocal microscopy image after BODIPY493/503 staining. N) Quantification of lipid droplet diameter M. O) Graphical conclusions for Fatty Acid Transfer. For all data, *in vitro*, n = 9, including three biological replicates and three technical replicates per biological replicate. Statistical significance was defined as a p-value < 0.05. Significance levels were denoted as follows: * P<0.05, ** P<0.01, *** P<0.001, and **** P<0.0001. Non-significant differences were labeled "ns".

Furthermore, upon *mfn2* knockdown, we unexpectedly found that HSL-mediated lipolysis was substantially suppressed, demonstrated by reduced hydrolase activity (Figure 5G-L) and decreased rates of LD hydrolysis (Figure 5M-N). Given that LD hydrolysis typically serves as the upstream process supplying fatty acids for mitochondrial *β*-oxidation, this result suggests that Mfn2-HSL-mediated tethering bidirectionally regulates the functions of both organelles. Collectively, these data demonstrate that Mfn2 and HSL jointly mediate a structural coupling between mitochondria and LDs that facilitates a rapid channel for efficient substrate transfer from LD hydrolysis to mitochondrial fatty acid *β*-oxidation (Figure 5O).

### 2.6. Mfn2 interacts with HSL through its GTPase domain and the N-terminal domain of HSL to maintain mitochondria-lipid droplet coupling

To elucidate the molecular mechanism by which Mfn2 and HSL mediate mitochondria-LD coupling, we first confirmed a direct protein interaction between the mitochondrial membrane protein Mfn2 and the LD-associated protein HSL. Their co-localization assays demonstrated a specific Mfn2-HSL interaction, which was significantly enhanced under creatine treatment (Figure 6A and S4A, B). To identify the precise interaction sites between Mfn2 and HSL, we separately isolated mitochondrial membrane and lipid droplet membrane fractions and performed co-immunoprecipitation (co-IP) assays (Figure 6B). The results demonstrated that Mfn2 interacted with HSL at the contact sites between mitochondria and lipid droplet membranes, and this interaction was significantly enhanced under creatine treatment (Figure 6C). We then conducted co-IP assays using Mfn2 as the bait protein in the purified PDM fraction. Results clearly demonstrated that the HSL isoform interacting with Mfn2 in the PDM fraction was predominantly phosphorylated HSL, with barely any non-phosphorylated inactive HSL detected in this Mfn2-binding complex (Figure 6D). To further verify this finding, we treated hepatocytes with a specific HSL phosphorylation inhibitor to block the phosphorylation of HSL, and repeated the co-IP assay in the purified PDM fraction. The results showed that inhibition of HSL phosphorylation significantly suppressed the physical interaction between Mfn2 and HSL on PDM (Figure 6E). Using molecular docking predictions combined with domain deletion experiments, we identified that Mfn2 primarily binds to HSL *via* its GTPase domain (amino acids 93-342) and the N-terminal domain of HSL (amino acids 7-317) (Figure 6F-H). This binding mode was further validated by co-localization microscopy in domain-deletion cellular models (Figure 6I, J and S4C, D). Importantly, deletion of either the GTPase domain of Mfn2 or the N-terminal domain of HSL markedly disrupted the physical coupling between mitochondria and LDs, as revealed by confocal analysis (Figure 6K-M). Taken together, these results establish that the direct interaction between the GTPase domain of mitochondrial Mfn2 and the N-terminal domain of LD-anchored HSL constitutes a key molecular mechanism underlying the maintenance of mitochondria-LD tethering.

**Fig. 6.**
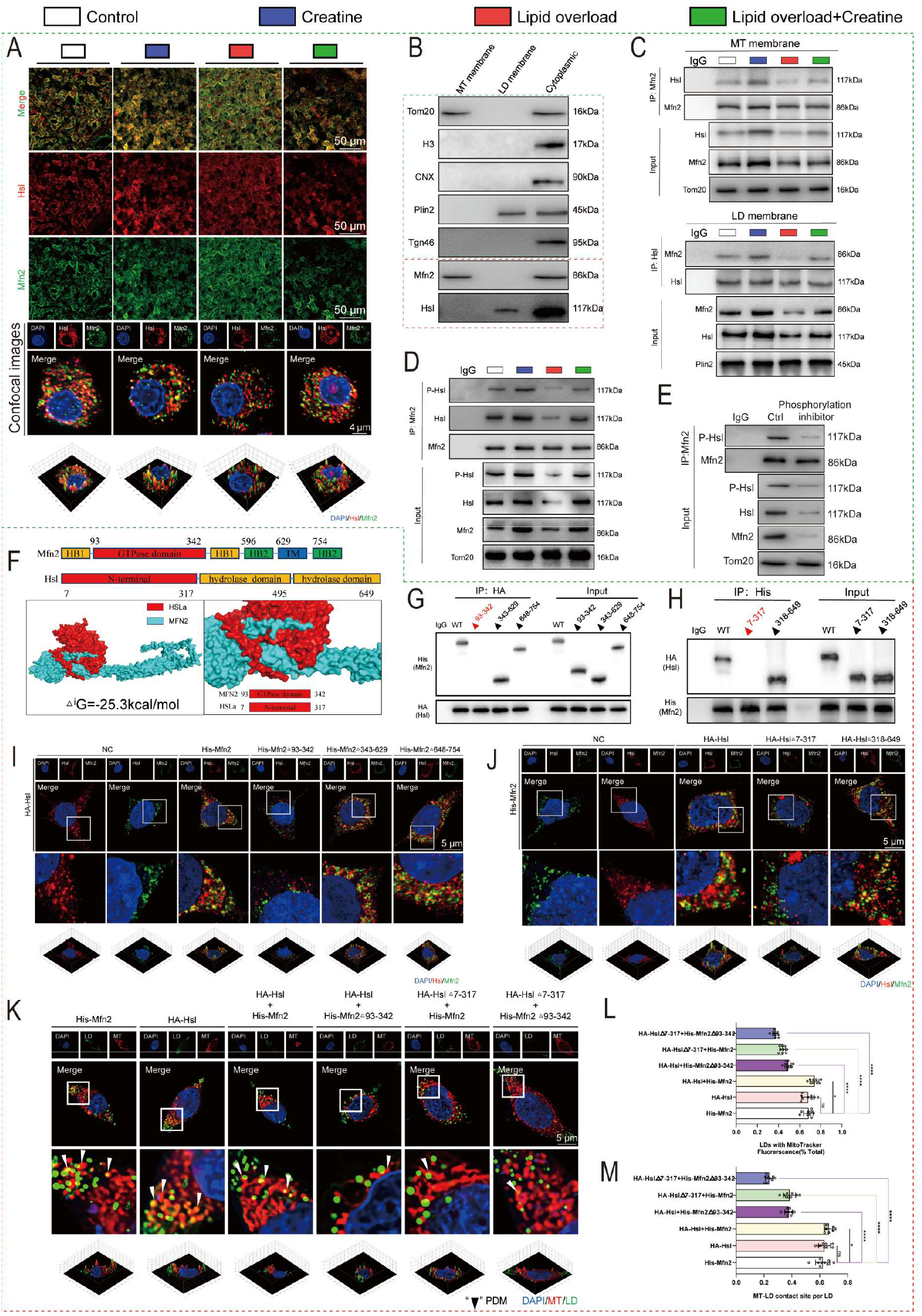
Mfn2 interacts with HSL through its GTPase domain and the N-terminal domain of HSL to maintain mitochondria-lipid droplet coupling. A) Representative confocal microscopic image of Mfn2 (green) and HSL (red) in the liver tissue section (*in vivo* experiments) and in yellow catfish hepatocytes (*in vitro* experiments). B) Western blot analysis of Tom20, H3, CNX, Plin2, Tgn46, Mfn2 and Hsl (*in vitro* experiments) C) CoIP of Mfn2 with HSL from MT membrane and LD membrane (*in vitro* experiments). D) CoIP of Mfn2 with HSL from PDM (*in vitro* experiments). E) CoIP of Mfn2 and p-HSL from PDM following phosphorylation inhibitor treatment (*in vitro* experiments). F) Protein docking prediction for Mfn2/HSL interaction and its core binding domain. G, H) CoIP of Mfn2 with HSL from 293T cells (which were transfected with wild-type Mfn2 and HSL, and their truncated mutants, *in vitro* experiments). I, J) Representative confocal microscopic image of Mfn2 with HSL from 293T cells (which were transfected with wild-type Mfn2 and HSL, and their truncated mutants, *in vitro* experiments). K) Representative confocal microscopy image after MitoTracker Deep Red and BODIPY493/503 staining (*in vitro* experiments). L) The fusion rate between MitoTracker and LDs in panel K. M) Mitochondria in contact with lipid droplets were quantified by % lipid droplet perimeter in panne K. For western blots, n=3 three biological replicates of one treatment. For the other data, *in vivo*, n = 9, 3 tanks for per diet with 3 fish randomly from each tank; *in vitro*, n = 9, including three biological replicates and three technical replicates per biological replicate. Statistical significance was defined as a p-value < 0.05. Significance levels were denoted as follows: * P<0.05, ** P<0.01, *** P<0.001, and **** P<0.0001. Non-significant differences were labeled "ns".

### 2.7. The precise molecular mechanisms for Mfn2-Hsl-mediated mitochondria-lipid droplet coupling and its regulatory role in LD hydrolysis

After identifying the interacting domains between Mfn2 and HSL, we further investigated the precise binding sites and their functional relevance in regulating LD hydrolysis. Through molecular docking predictions followed by site-directed mutagenesis and co-immunoprecipitation assays, we identified specific residues— Mfn2 (Lys192) and HSL (Thr203)—as critical for their direct interaction (Figure 7A, B). This finding was corroborated by impaired co-localization between Mfn2 and HSL in cells expressing Mfn2 (K192R) or HSL (T203S) mutants (Figure 7C, D and S5A, B). Importantly, disruption of either residue severely compromised mitochondria-LD coupling, providing precise molecular targets for this inter-organelle tethering (Figure 7E-G).

**Fig. 7.**
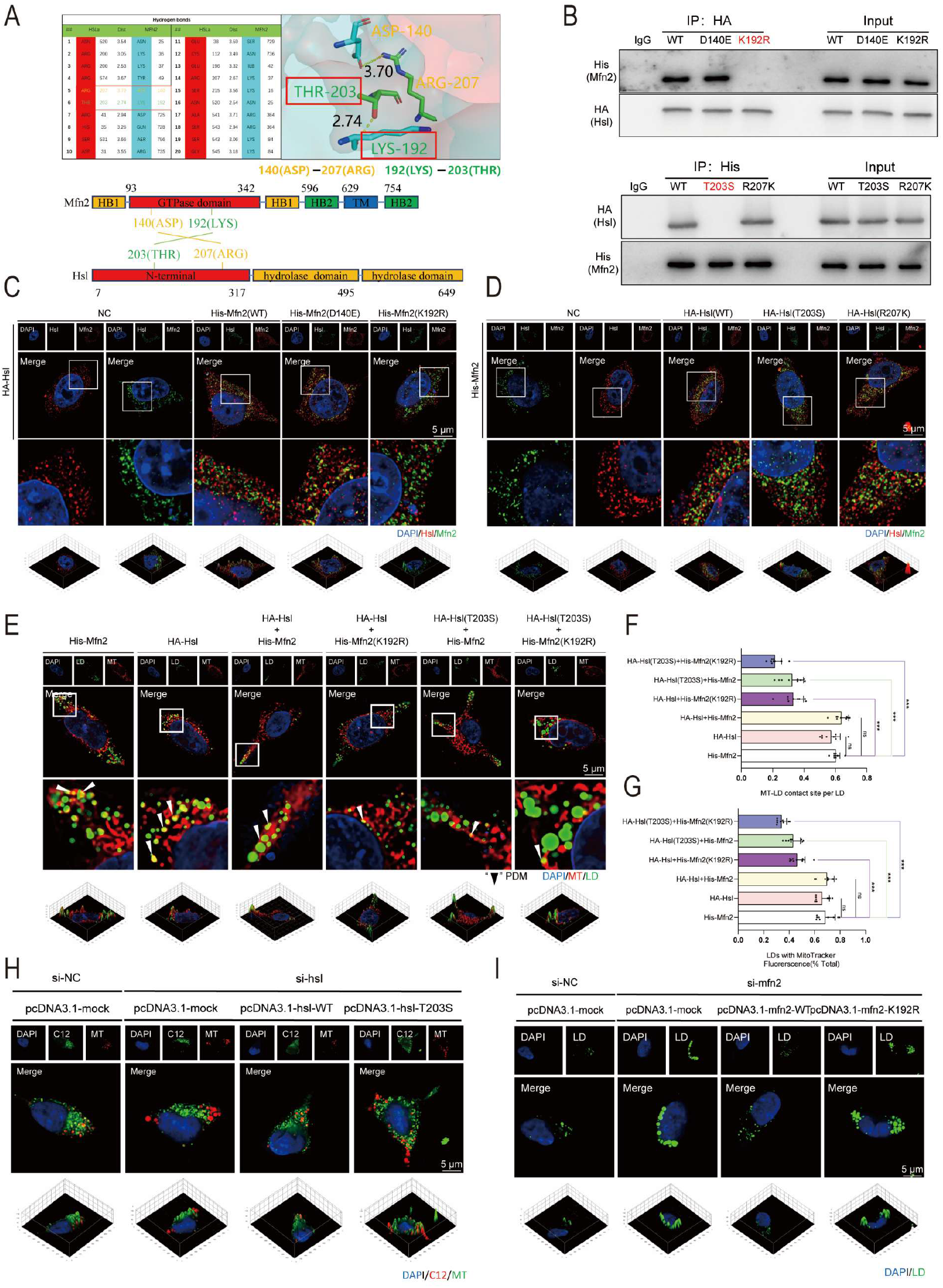
The precise molecular mechanisms for Mfn2-HSL-mediated mitochondria-lipid droplet coupling. (*in vitro* experiments). A) Protein docking prediction for Mfn2/HSL interaction and its core binding site. B) CoIP of Mfn2 with HSL from 293T cells (which were transfected with wild-type Mfn2 and HSL, and their point mutations). C, D) Representative confocal microscopic image of Mfn2 with HSL from 293T cells (which were transfected with wild-type Mfn2 and HSL, and their point mutants). E) Representative confocal microscopy image after MitoTracker Deep Red and BODIPY493/503 staining. F) Mitochondria in contact with lipid droplets were quantified by % lipid droplet perimeter in panel E. G) The fusion rate between MitoTracker and LDs in panel E. H) Representative confocal images of mitochondria and Red C12 480/508, scale bars, 5µm. I) Representative confocal microscopy image after BODIPY493/503 staining scale bars, 5µm. For the other data, *in vitro*, n = 9, including three biological replicates and three technical replicates per biological replicate. Statistical significance was defined as a p-value < 0.05. Significance levels were denoted as follows: * P<0.05, ** P<0.01, *** P<0.001, and **** P<0.0001. Non-significant differences were labeled "ns".

To investigate the effect of this interaction site on biological function, we performed a functional rescue experiment: endogenous *hsl* was first knocked down, followed by re-expression of an *hsl* mutant that only disrupted the interaction between HSL and Mfn2. The mutant site was designed according to the point mutations presented in Figure 6. The results showed that re-expression of this mutant significantly attenuated the trafficking of C12-labeled fatty acids into mitochondria (Figure 7H and S5C) and still led to a significant decrease in mitochondrial fatty acid *β*-oxidation (Figure S5D, E). Furthermore, the point mutation of Mfn2 (Lys192) significantly suppressed HSL activity and reduced the levels of related metabolic intermediates (Figure S5F-I). Consistently, confocal microscopy confirmed inhibited lipid droplet lipolysis (Figure 7I and S5J), indicating that the Mfn2 (Lys192)-HSL (Thr203) interaction not only maintains structural coupling but also regulates lipolytic processes and fatty acid *β*-oxidation. These results demonstrate that the Mfn2-HSL interaction at these specific residues is essential for both the physical tethering and the functional coordination between mitochondrial fatty acid *β*-oxidation and LD hydrolysis.

### 2.8. Conservation of Mfn2-HSL-mediated mitochondria-lipid droplet coupling and its regulatory role in LD hydrolysis and mitochondrial fatty acid *β*-oxidation

Given the widespread presence of mitochondria-LD interactions across vertebrates,^25^ we investigated whether the regulatory role of Mfn2-HSL in maintaining this inter-organelle tethering and its associated metabolic functions is evolutionarily conserved. We systematically evaluated the effects of *mfn2* or *hsl* knockdown in hepatocyte models from multiple vertebrate species, including humans, mice, chickens and frogs. Under lipid overload and creatine treatment conditions, all species exhibited phenotypes consistent with those observed in fish: creatine attenuated lipid overload-induced mitochondrial fragmentation and enhanced both LD hydrolysis and subsequent mitochondrial fatty acid *β*-oxidation (Figure S6A-D). Furthermore, either *mfn2* or *hsl* knockdown disrupted mitochondria-LD coupling in each species, indicating a conserved structural role of the Mfn2-HSL complex in maintaining inter-organelle tethering (Figure 8A1-D1, A2-D2). Importantly, *mfn2* knockdown consistently suppressed LD hydrolysis (Figure 8A3-A6, B3-B6, C3-C6, D3-D6), and *hsl* knockdown consistently inhibited fatty acid oxidation across species (Figure 8A7-A10, B7-B10, C7-C10, D7-D10), demonstrating that the Mfn2-HSL-mediated coupling not only maintains physical contact but also exerts a conserved regulatory influence on hepatic lipid metabolism, particularly in coordinating lipolysis with mitochondrial fatty acid *β*-oxidation.

**Fig. 8.**
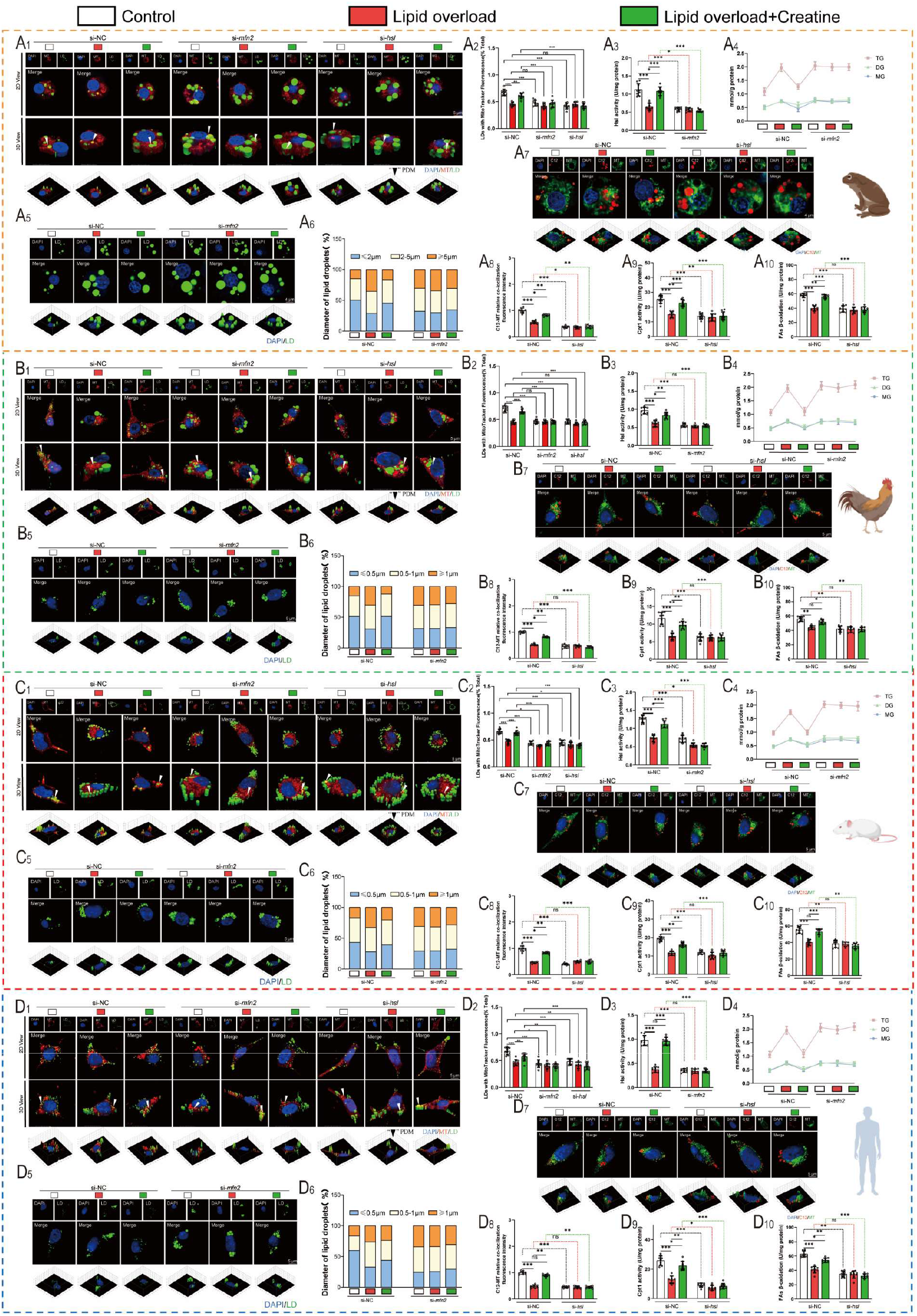
Conservation of Mfn2-HSL-mediated mitochondria-lipid droplet coupling and its regulatory role in LD hydrolysis and mitochondrial fatty acid *β*-oxidation (*in vitro* experiments). A_1_) Representative confocal microscopy image after MitoTracker Deep Red and BODIPY493/503 staining in hepatocytes from frog, scale bars, 5µm. A_2_) The fusion rate between MitoTracker and LDs in panel A_1_. A_3_) HSL activity in hepatocytes from frog. A_4_) TG, DG and MG content in hepatocytes from frog. A_5_) Representative confocal microscopy image after BODIPY493/503 and Red C12 480/508 staining, scale bars, 4µm. A_6_) Quantification of lipid droplet diameter in panel A_5_. A_7_) Representative confocal images of mitochondria and Red C12 480/508, scale bars, 4µm. A_8_) The fusion rate between mitochondria and Red C12 in panel A_7_. A_9_) Cpt1 activity of isolated mitochondria from cells. A_10_) FAs *β*-oxidation rate of isolated mitochondria from cells. B_1_-B_10_, C_1_-C_10_ and D_1_-D_10_): The conserved mechanism by which creatine alleviates lipid deposition in hepatocytes from chicken (LMH), mice (NCTC), and human (HepG2), respectively. For all data, *in vitro*, n = 9, including three biological replicates and three technical replicates per biological replicate. Statistical significance was defined as a p-value < 0.05. Significance levels were denoted as follows: * P<0.05, ** P<0.01, *** P<0.001, and **** P<0.0001. Non-significant differences were labeled "ns"

## 3. DISCUSSION

The physical coupling between mitochondria and lipid droplets (LDs) shortens the transport distance of fatty acids from LDs to mitochondria,^5^ theoretically creating an efficient metabolic channel for lipolysis and fatty acid oxidation. We demonstrate for the first time that Mfn2/HSL-mediated mitochondria-LD tethering underpins this rapid metabolic channel: Mfn2 activates HSL hydrolytic activity through direct protein-protein interaction sites, thereby accelerating the LDs release of free fatty acids for mitochondrial *β*-oxidation. Additionally, we found that dietary creatine significantly enhances lipolysis and oxidation by reinforcing this coupling, thereby alleviating hepatic lipid accumulation. This mechanism is highly conserved in vertebrates, highlighting its potential as a universal metabolic target. Collectively, our findings propose a therapeutic strategy centered on strengthening mitochondria-LD coupling to optimize fatty acid catabolism efficiency, offering promising avenues for the treatment of hepatic steatosis.

To explore the physiological relevance of the mitochondria-LD coupling mechanism, we investigated its response to excessive hepatic lipid deposition using a high-fat diet (Lipid overload) model in yellow catfish. The present study demonstrates that Lipid overload significantly disrupts the mitochondria-LD coupling structure in hepatocytes—a finding consistent with our previous observations.^12^ Creatine has shown potential in reducing hepatic lipid deposition and improving mitochondrial function.^13,15^ In this present study, dietary creatine supplementation restored this structural integrity, thereby promoting the formation of peridroplet mitochondria (PDM). Under conditions where total mitochondrial content remained constant, fused mitochondria exhibited a higher propensity for PDM formation compared to fragmented mitochondria. Beyond further validating our earlier conclusions,^26^ this result also aligns with the research outcomes of Freyre.^1^ Mechanistically, creatine specifically activated the *hnf4α*-responsive element within the *mfn2* gene promoter region, significantly enhancing *mfn2* transcriptional activity. This activation, in turn, promoted mitochondrial fusion and subsequent PDM formation. Subsequently, site-directed mutagenesis of the endogenous *hnf4α* binding element completely abrogated the promotive effects of creatine on mitochondrial fusion and the formation of lipid droplet-mitochondria contact sites. Meanwhile, both lipid droplet lipolysis and mitochondrial fatty acid *β*-oxidation were significantly reduced. The above observations are consistent with the conclusions reported by Xu ^27^ who confirmed that *hnf4α* acts as a core transcriptional regulator of lipolysis and mitochondrial fatty acid *β*-oxidation.

Previous studies have demonstrated that PDM facilitate lipid droplet expansion and exhibit a limited capacity for fatty acid oxidation.^17^ Notably, that study was exclusively conducted in adipocytes. Herein, our findings verify that in hepatocytes, PDM harbor substantially higher activity of fatty acid *β*-oxidation than cytosolic mitochondria, which is consistent with previous related reports.^28,29^ We therefore propose that PDM may exert distinct functions in a cell-type-specific manner. Then through comprehensive analysis of hepatic lipid metabolic parameters, we confirmed that dietary creatine effectively attenuates Lipid overload-induced hepatic lipid accumulation, in accordance with earlier observation.^15,30–32^ Intriguingly, the present study also indicated dietary creatine exerts this protective effect by coordinately activating lipolysis and fatty acid *β*-oxidation, which corroborates the observed enhancement in *β*-oxidation efficiency of PDM.

Having demonstrated that dietary creatine enhances mitochondria-LD coupling through Mfn2 and activates lipolysis and *β*-oxidation, we next aimed to elucidate the underlying molecular mechanism. Lipid catabolism typically proceeds through a well-characterized sequence: stored lipids (predominantly in LDs) are hydrolyzed to release fatty acids, which are then shuttled to mitochondria for subsequent oxidation.^33^ Concomitant with the severe disruption of mitochondria-LD coupling induced by si-*mfn2*, our data revealed that si-*mfn2* also substantially attenuated hepatic LD lipolysis. This finding suggests that Mfn2-mediated mitochondria-LD coupling directly influences upstream LD lipolysis processes, which is consistent with the findings of Hu.^10^ The hydrolysis of stored fats is generally mediated by a cascade of lipases.^34–36^ Among these, si-*mfn2* exerted the most pronounced inhibitory effect on HSL activity, indicating that HSL plays a pivotal role in Mfn2-regulated lipolysis. First, si-*hsl* similarly disrupted mitochondria-LD coupling structures, and impaired Mfn2-dependent mitochondrial fusion. Additionally, si-*hsl* significantly reduced *β*-oxidation efficiency, likely due to an enlarged spatial gap between LDs and mitochondria that impedes fatty acid shuttling—a known rate-limiting step for lipid lipolysis and oxidation.^4^ Collectively, these results unveil a bidirectional regulatory loop: Mfn2 maintains mitochondria-LD coupling to facilitate HSL-dependent lipolysis, while HSL, in turn, sustains the integrity of this coupling and ensures efficient fatty acid *β*-oxidation. Notably, our study identifies that creatine exerts its beneficial effects on hepatic lipid metabolism through this regulatory loop. These findings highlight a sophisticated spatial metabolic crosstalk between mitochondria and LDs, which enables rapid hepatic lipid hydrolysis and *β*-oxidation.

Mfn2 localizes to the mitochondrial outer membrane,^37^ while HSL translocates to the surface of lipid droplets after phosphorylation.^38^ These proteins exhibit distinct subcellular distributions has been previously validated in our studies on yellow catfish.^12,39^ Building on this foundation, the present study advances our understanding by elucidating the precise molecular mechanisms through which Mfn2/HSL regulate mitochondria-LD coupling structures and their associated lipid metabolism functions. Our experimental results confirm a direct protein-protein interaction between Mfn2 and p-HSL. Specifically, Mfn2 binds to HSL *via* its GTP domain, which interacts with the N-terminal of HSL. Previous studies have indicated that the GTP domain of Mfn2 plays an important regulatory role in its GTP hydrolysis function,^40,41^ while the N-terminal of HSL is a potential region for protein-protein interactions.^42^ In this present study, critical functional analyses revealed that deletion of either the GTP domain in Mfn2 or the N-terminal in HSL substantially impaired mitochondria-LD coupling, definitively establishing the essential role of the Mfn2/HSL interaction in maintaining this coupling. Furthermore, we precisely mapped the key interaction sites to Mfn2-192lys and HSL-203thr. Interestingly, mutation analysis demonstrated that Mfn2-192lys regulates HSL enzymatic activity, as its mutation significantly attenuated HSL-mediated lipid droplet hydrolysis. This suggests that Mfn2 modulates upstream lipolysis through this specific residue. This study provides the first molecular evidence that Mfn2 promotes LD hydrolysis *via* mediating mitochondria-lipid droplet coupling. Importantly, we identified that creatine supplementation can enhance metabolic coordination between upstream lipolysis and downstream mitochondrial fatty acid oxidation *via* Mfn2-HSL-mediated mitochondria-LD coupling, thereby establishing an efficient lipolysis and *β*-oxidation metabolic axis and alleviating the excessive hepatic lipid deposition These findings highlight the interaction between Mfn2 and p-HSL as a promising therapeutic target for ameliorating lipid overload-induced hepatic steatosis by restoring mitochondria-LD coupling and optimizing lipid metabolic flux.

Given the universal presence of mitochondria and LDs in vertebrates,^25^ along with the high evolutionary conservation of Mfn2 and HSL across species,^19,20^ another key finding of this study is the first discovery and confirmation that the Mfn2/HSL-regulated mitochondria-LD coupling structure, together with its regulatory mechanism in hepatic lipid metabolism, is evolutionarily conserved among vertebrates. Specifically, we demonstrated that inhibiting either Mfn2 or HSL disrupts mitochondria-LD coupling in all tested vertebrate cells. Functionally, consistent with observations in yellow catfish, si-*mfn2* significantly reduced HSL activity and suppressed LD hydrolysis; in contrast, HSL interference also affect Mfn2-mediated mitochondrial fusion and markedly impaired fatty acid oxidation in hepatocytes—also likely due to the physical dissociation of mitochondria from LDs. Collectively, these findings support the conclusion that vertebrates universally possess an Mfn2/HSL-dependent mitochondria-LD coupling structure, which acts as a critical pathway for rapid LD breakdown and mitochondrial fatty acid *β*-oxidation.

Collectively, this study is the first to clarify that mitochondria-LD coupling mediated by Mfn2/HSL establishes an efficient metabolic axis that regulates LD lipolysis and mitochondrial *β*-oxidation. Key discoveries include: (1) Mfn2 directly interacts with HSL *via* specific binding domains (Mfn2-192lys / HSL-203thr), with Mfn2-192lys critically regulating HSL hydrolytic activity; (2) Creatine enhances this coupling by activating *hnf4α*-dependent Mfn2 transcription, promoting PDM formation and superior *β*-oxidation capacity; (3) A bidirectional regulatory loop exists wherein Mfn2 sustains HSL-dependent lipolysis, while HSL maintains coupling integrity for efficient fatty acid shuttling. Notably, this Mfn2/HSL mechanism is evolutionarily conserved across vertebrates, highlighting its potential as a universal therapeutic target for hepatic steatosis. Our findings propose strengthening mitochondria-LD coupling as a strategy to optimize hepatic lipid catabolism, offering novel insights into spatiotemporal metabolic coordination. A working model of aforementioned mechanism is shown in Figure 9.

**Fig. 9.**
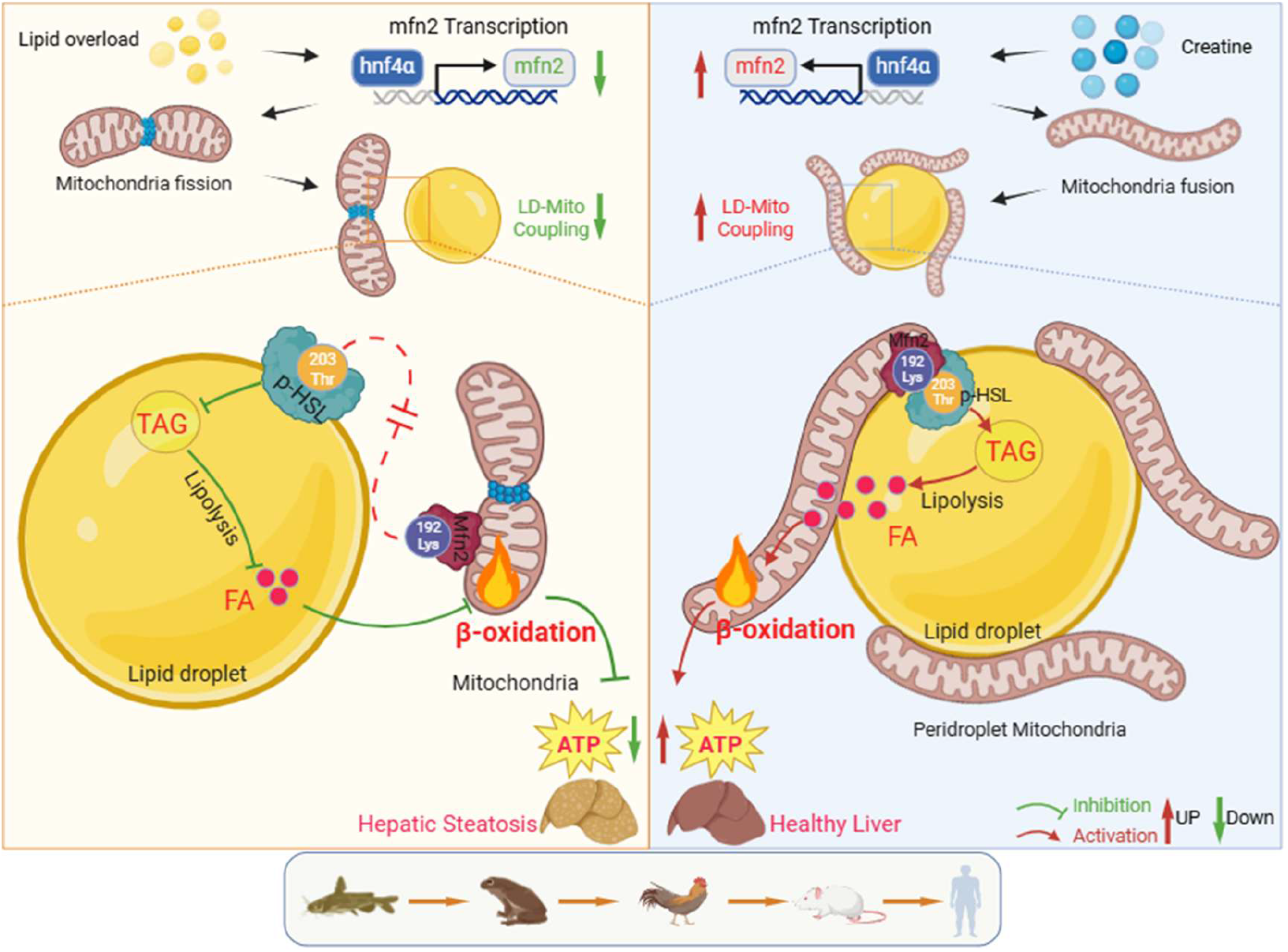
Graphical conclusions of a novel and evolutionarily conserved metabolic axis of Mfn2-HSL-mediated mitochondria-lipid droplet coupling for efficient hepatic lipid catabolism.

## 4. EXPERIMENTAL MODEL AND STUDY PARTICIPANT DETAILS

### Ethical statement

Huazhong Agricultural University’s (HZAU) institutional ethical guidelines for the care and use of laboratory animals were followed throughout all investigations, and were approved by the Ethical Committee of HZAU (Approval No.: Fish-2024-0127).

### Experiment. 1, Animals feeding and sampling (in vivo experiment)

The experimental diet formulation for yellow catfish was adapted from our previous publications.^43^ Four dietary treatments were prepared with different lipid levels and creatine supplementation (Table S2): 10.6% lipid and 0 g·kg⁻¹ creatine (group 1: control), 10.6% lipid and 20 g·kg⁻¹ creatine (group 2: Cr), 14.4% lipid and 0 g·kg⁻¹ creatine (group 3: lipid overload), and 14.4% lipid and 20 g·kg⁻¹ creatine (group 4: lipid overload + Cr). A 1:1 (w/w) mixture of fish oil and soybean oil served as the lipid source. At the start of the experiment, 360 fish of uniform size (5.21 ± 0.03 g, mean ± SEM) were selected and randomly distributed into 12 tanks (300 L) in a completely randomized design, with three replicates per diet and 30 fish per tank. The fish were fed twice daily at 08:00 and 16:00 for 8 weeks. Throughout the trial, water quality was maintained as follows: dissolved oxygen 6.43 ± 0.18 mg/L, temperature 28.3 ± 0.3°C, pH 7.79 ± 0.14, and NH₄-N 0.049 ± 0.002 mg/L.

At the end of the feeding trial, fish were fasted for 24 h and euthanized with buffered MS-222 (200 mg/L). Surviving fish in each tank were counted and bulk-weighed for the calculation of survival, weight gain (WG), specific growth rate (SGR), and feed conversion ratio (FCR). Growth and morphological data are summarized in Table S1. For each diet group, liver tissues from nine fish across three tanks (3 fish per tank) were sampled for histology, ultrastructure, and immunofluorescence observations. The remaining liver samples were snap-frozen in liquid nitrogen and stored at -80°C for subsequent assays, including triglyceride content, enzyme activity, PeriDroplet Mitochondria (PDM) extraction, and gene and protein expression analyses.

### Experiment. 2, Culture and treatments of hepatocyte lines or primary hepatocytes from varied vertebrates (in vitro experiment)

Primary hepatocyte culture and treatment: yellow catfish and frog were obtained from the Laboratory Animal Center of Huazhong Agricultural University. Primary hepatocytes were isolated and cultured as described previously.^44^ The cells were incubated at 22 °C or 28 °C. Hepatocellular cell line culture and treatment: we employed multiple liver cell lines including HepG2 (human hepatocellular carcinoma), HEK-293T (human embryonic kidney), NCTC (mouse normal liver cells), and LMH (chicken hepatocellular carcinoma) for cross-species validation. Based on previous studies and cell viability assays,^45^ we treated cells with a fatty acid mixture (1mM, PA: OA = 1:1) and creatine for 48 h (Figure S7). All treatments were performed in triplicate, with each replicate consisting of pooled hepatocytes from three individuals.

### Sample analysis

#### Oil red O, Hematoxylin and Eosin (H&E), Bodipy 493/503 staining, MitoTracker Staining and transmission electronic microscopy (TEM) observation

For histological observation, sagittal sections with a thickness of 6-8 µm were prepared and stained with H&E or Oil red O for light microscopy, as previously described.^38^ Staining of lipid droplets with Bodipy 493/503 (5μg/ml) and mitochondria with MitoTracker Deep Red FM 644/665 (100 nm) was performed according to our established protocols.^46^ Briefly, after 48 h of treatment, cells were washed with PBS, incubated with Bodipy 493/503 or MitoTracker Deep Red FM 644/665 for 30 min and imaged by laser scanning confocal microscopy. All image assessments were performed in a double-blind manner. Ten fields per sample were quantified using ImageJ software to measure the relative area of LDs and mitochondrion in hepatocyte or Oil Red O staining and the relative area of vacuoles in H&E staining, as previously described.^23^ Additionally, hepatocyte ultrastructure was examined by TEM using methodologies detailed in our prior work.^47^ Ten images were acquired from each liver sample, and ten fields per sample were analyzed with ImageJ to determine the morphology and quantity of mitochondria. Mitochondria were classified into distinct morphological phenotypes based on their length: Elongated (≥1 μm), Intermediate (0.5-1 μm), and Fragmented (≤0.5 μm).

#### Quantification of creatine, triglyceride (TG), diacylglycerol (DG), monoglyceride (MG), and non-esterified fatty acids (NEFAs)

Creatine levels in yellow catfish liver tissues were quantified using commercial kit (Abcam Cat: ab65339) following manufacturer’s instructions. The measurement of TG, DG and MG contents was performed following our established protocols,^12^ using TG Assay Kit (A110-1-1, Nanjing Jiancheng Bioengineering Institute), DG and MG Assay Kit (YX-130107F and YX-040700F, Sinobestbio, China). In brief, liver tissues were homogenized and centrifuged at 12,000 rpm for 5 minutes. Subsequently, 100 µL of the supernatant was transferred to a 24-well plate and mixed with the corresponding detection reagents. Luminescent signals were measured using a microplate reader (BMG Labtech, Offenburg, Germany). The protein concentration was measured using the commercial kits (Nanjing Jiancheng Bioengineering Institute, Nanjing, China; A045-2-2). The contents of NEFAs in the liver or hepatocytes were evaluated using tissue NEFAs assay kits (Nanjing Jiancheng Bioengineering Institute, Nanjing, China; A042-2-1), following the manufacturer’s instructions.

#### Assay the activities of adipose triglyceride lipase (ATGL), hormone-sensitive lipase (HSL), and monoacylglycerol lipase (MGL)

The activities of ATGL, HSL and MGL were assayed following the method of Schweiger.^48^ The substrates used were: 1.67 mM unlabeled triolein (Sigma, CAS No. 122-32-7) with 10μ Ci/ml [carboxyl-¹⁴C]triolein; 0.3 mM unlabeled diolein with 10μ Ci/ml 1-¹⁴C-labeled diolein (1,3-dioleoylglycerol [1-¹⁴C]oleoyl); or 0.3 mM unlabeled monoolein (Sigma, CAS No. 111-03-5) with 10μ Ci/ml 1-¹⁴C-labeled monoolein (1-monooleoylglycerol [1-¹⁴C]oleoyl). Briefly, tissues were homogenized in buffer (50 mM Tris [pH 7.4], 0.1 M sucrose, 1 mM EDTA). After homogenization, samples were incubated with substrates at 37 °C for 1 h. Reactions were stopped by adding 3.5 ml methanol-chloroform-heptane (10:9:7). Fatty acids were extracted with 1 ml of 0.1 M potassium carbonate-0.1 M boric acid buffer (pH 10.5), and radioactivity in the upper phase was measured by scintillation counting.

#### Assay of Cpt1 Activities, ATP content and Mitochondrial Fatty Acid *β*-Oxidation

The Cpt1 activity assay was determined according to our publication.^38^ The enzymatic assay, based on the method of Fiol,^49^ quantifies initial CoA-SH generation *via* the 5,5′-dithio-bis-(2-nitrobenzoic acid) (DTNB) reaction, wherein palmitoyl-CoA is converted in the presence of L-carnitine, with absorbance monitored at 412 nm. One international unit (IU) of Cpt1 activity was defined as the production of 1 μmol of product per minute per mg of mitochondrial protein at 25°C. ATP was measured by using an ATP Colorimetric/Fluorometric Assay Kit (S0026; Beyotime, Nantong, China) according to the manufacturer’s instructions. Mitochondrial fatty acid *β*-oxidation rate was determined as previously described.^50^ Briefly, *β*-oxidation was assessed using 100 mM PA (supplemented with 1 μ Ci [1-14C]-PA, specific activity 60 μ Ci/ mmol, NEC075H050UC, PerkinElmer, Pittsburgh, PA, USA) as the radiolabeled substrate.

### RNA Extraction and Real-Time Quantitative Polymerase Chain Reaction(qPCR)

Total RNA was isolated using Trizol reagent, and then cDNA was synthesized using a reverse transcription kit. Gene transcription level analysis was performed by real-time quantitative PCR (qPCR), following our previously published study.^51^ The primer sequences used in this study are detailed in Table S3-S7. Transcriptional stability of nine housekeeping genes (*β-actin, tbp, elfa, b2m, hprt, ubce, tuba, gapdh, 18s rRNA*) was tested, revealing *gapdh* and *β-actin* (M = 0.25) as most stably expressed across all experimental conditions, thus selected as endogenous controls. Relative expression was calculated *via* the 2−ΔΔCt method.

### DNA Isolation and qPCR for mtDNA

According to our previously published study,^39^ the mtDNA was isolated by using a commercial kit (K280-50; MT DNA Isolation Kit; Biovision, Mountain View, CA, USA). mtDNA copy number was determined using the mitochondrial *nd2* gene, with 18S rRNA as the reference standard. The primer sequence is detailed in Table S8. Convert cytoplasmic mitochondrial DNA concentration to copy number by DNA copy number calculator, and all samples were assayed in parallel with standards.

### Protein docking for Mfn2 and HSL

The protein structure of yellow catfish was modeled using the SWISS-MODEL server, with crystal structure data sourced from the PDB database (Mfn2 PDB ID: 6jfm; HSL PDB ID: 8zvq). Suitable templates were selected from these data. The ZDOCK module in Discovery Studio 4.0 was employed to perform Mfn2-HSL docking, and the best docking conformation was selected from the generated models using PyMOL.

### Cell Transfection and Plasmid Construction

To explore the functions of Mfn2 and HSL in PDM-regulated hepatic lipid metabolism, species-specific siRNAs targeting *mfn2* and *hsl* were designed. Following our previous protocol,^52^ 50 nM siRNA per well was transfected into primary yellow catfish and frog hepatocytes using Entranster™-R4000 reagent (Engreen, China). Target siRNA sequences are listed in Table S9 and were synthesized by GenePharma (Shanghai, China). Knockdown efficiency was validated at mRNA and protein levels (Figure S8). For cross-species validation, corresponding siRNAs targeting *mfn2* and *hsl* (specific to chicken, mice, or human) were transfected into LMH, NCTC, and HepG2 cell lines. For mapping protein interaction sites, site-directed mutants of Mfn2 (D140E, K192R) and HSL (T203S, R207K) were generated using the Mut Express II Fast Mutagenesis Kit (Vazyme). We inserted three categories of ORFs for each gene into His-tagged or HA-tagged pcDNA3.1(+) vectors: full-length ORFs, interaction-sequence deletion variants (Mfn2 Δ93-342, Δ343-629, Δ648-754; HSL Δ7-317, Δ318-649), and point-mutation variants (Mfn2 D140E, K192R; HSL T203S, R207K). The primer sequences are listed in Table S10. Plasmids were transiently transfected into HEK-293T cells using Lipofectamine 2000 (Invitrogen).

### Western blot and immunoprecipitation

Based on our previous study,^53^ western blotting was used to detect the protein expression levels of HSL, Cpt1, Opa1, Mfn1, Mfn2, Tom20 and Gapdh. Briefly, liver samples and cell lysates were homogenized in RIPA buffer (Thermo Fisher) with protease inhibitor (Sigma-Aldrich). Protein concentration was determined using a BCA kit (Nanjing Jiancheng), and 15 μg protein per lane was separated by SDS-PAGE (10%, 12% or 15% gels) and transferred to PVDF membranes (Millipore). After blocking with 8% nonfat milk, membranes were incubated with primary antibodies at 4°C overnight, then with HRP-conjugated secondary antibody at room temperature for 2 h. Protein bands were detected using an Odyssey Imager (Li-Cor) with ECL (Bio-Rad) and quantified *via* Image Lab software, normalized to Gapdh.

For immunoprecipitation, cells were lysed in protein lysis buffer supplemented with protease inhibitors. Lysates were incubated overnight at 4 °C with anti-Mfn2 (Cat No. 12186-1-AP; Proteintech), anti-HSL (Cat No. 17333-1-AP; Proteintech), anti-p-HSL (Cat No. 86398-1-RR; Proteintech), anti-His (Cat No. 66005-1-Ig; Proteintech), or anti-Flag (Cat No. 20543-1-AP; Proteintech) antibodies as bait. Protein A/G agarose was added and incubated for 7 h, followed by five washes with PBS containing PMSF. The immunoprecipitated complexes were then analyzed by Western blot using the corresponding antibodies as prey.

### Composition of Mfn2-protein complexes formed in liver lysates identified by LC-MS/MS analysis

According to our previously described protocol,^12^ the composition of the Mfn2-protein complexes was determined. LC-MS/MS analysis was conducted as previously described.^45^ Cells were lysed in protein lysis buffer supplemented with protease inhibitors. Lysates were incubated overnight at 4 °C with anti-Mfn2 antibody as bait, followed by a 7-hour incubation with Protein A/G agarose. After washing with PBS containing PMSF, immunoprecipitated complexes were obtained. These complexes were incubated with SDS loading buffer and separated by electrophoresis. Electrophoresis was stopped when the bromophenol blue dye had migrated approximately 1.5 cm into the resolving gel. The protein-containing gel band was excised, reduced, alkylated, and subjected to in-gel tryptic digestion. The resulting peptides were separated on an UltiMate 3000 RSL Cnano system coupled online to a Q Exactive HF hybrid quadrupole-orbitrap mass spectrometer and analyzed. MS data were processed using MaxQuant software v1.6.6 against the yellow catfish NCBI database (*Pelteobagrus fulvidraco*, 20230407). Proteins were quantified by MaxQuant’s label-free quantification algorithm based on MS1 intensity, and the relative abundance of each protein in the corona was normalized to the total protein intensity.

### Immunofluorescence and 2D/or 3D confocal analysis

Immunofluorescence and confocal microscopy were performed to evaluate the colocalization of Mfn2 and HSL, also mitochondria and lipid droplets in hepatocytes, as previously described.^12^ For tissue sections: frozen liver sections were thawed, permeabilized with 0.1% Triton X-100, blocked with 5% FBS, and incubated with primary antibodies (anti-Mfn2, anti-HSL) at 4°C overnight, then with secondary antibodies at room temperature for 1 h, before mounting with an anti-fluorescence quencher. Nuclei and lipid droplets were stained with DAPI (5μg/ml) and Bodipy 493/503, respectively. For fatty acid transfer assays: cells were labeled with Bodipy C12 (1 mM) for 16 h and mitochondria with MitoTracker Deep Red. For subcellular localization: hepatocytes were fixed with 4% paraformaldehyde, permeabilized, blocked, and incubated with primary antibodies (anti-His, anti-Flag, anti-Mfn2, anti-HSL) and corresponding secondary antibodies; nuclei were stained with DAPI. Super-resolution imaging was performed on a Nikon N-SIM system. Images were processed using NIS-Elements, with intensity quantified in Image-Pro Plus and colocalization assessed in ImageJ. For 3D imaging: images were acquired with a Plan-Apochromat ×40 objective and triple zoom.

### Isolation of perilipid droplet mitochondria (PDM) and cytoplasmic mitochondria (CM)

PDM and CM isolation followed our previous study,^12^ with all procedures conducted on ice. Briefly, liver samples were homogenized in Sucrose-HEPES-EGTA buffer with BSA. Following centrifugation at 900×g for 11 min at 4°C, the upper fat layer and supernatant were transferred to fresh tubes, and this process was repeated several times. Samples were then resuspended in SHE+BSA and centrifuged at 9000×g for 18 min at 4°C. The final pellet was resuspended in SHE without BSA: the pellet derived from the fat layer was defined as PDM, and that from the supernatant as CM.

The *in vitro* co-incubation assay between lipid droplets and mitochondria was performed following a previously documented protocol established by Zervopoulos et al.^54^ Purified mitochondria and lipid droplets were separately resuspended in a commercial cytoplasmic extraction buffer (Cat. No. K266-100, Abcam). Lipid droplets containing 80 μg of total protein were thoroughly blended with an equivalent amount of mitochondria in 30 mm confocal glass-bottom dishes (P35G-1, MatTek). After incubation at 28 °C for 1 h, the samples were fixed with 2% paraformaldehyde prepared in 1× phosphate-buffered saline (PBS). Immunofluorescence staining was subsequently conducted in accordance with the experimental procedures described above.

### Promoter Cloning, Dual Luciferase Reporter Assay and Electrophoretic Mobility Shift Assay (EMSA)

The *mfn2* promoter from yellow catfish was cloned and inserted into the pGL3-Basic vector following a previously described method^51^ using the ClonExpress™ II One Step Cloning Kit (Vazyme, C112-01). Putative *hnf4α* binding sites in the mfn2 promoter were predicted using the JASPAR database. Site-directed mutagenesis of the *hnf4α* binding region was carried out with a Site-Directed Mutagenesis Kit (Vazyme, C216-01), using mutagenic primers provided in Table S11. Promoter activity was measured with the Dual-Luciferase Reporter Assay System. Binding of *hnf4*α to the mfn2 promoter was verified by electrophoretic mobility shift assay (EMSA). Nuclear proteins were extracted using a commercial kit (Beyotime, P0027). Reaction mixtures were incubated for 30 min as instructed in the LightShift™ Chemiluminescent EMSA Kit protocol (Thermo Fisher Scientific, 20148). Samples were separated on 6% native polyacrylamide gels, transferred to nylon membranes, and UV-crosslinked. Complex signals were detected by chemiluminescence. Competition assays were performed with a 200-fold excess of unlabeled oligonucleotide duplexes (wild-type or mutated).

### Statistical Analysis

All data were expressed as mean ± standard error of the mean (S.E.M.) for statistical analysis using GraphPad Prism software 10.0.2. For western blots, n=3 three biological replicates of one treatment. For the other data, *in vivo*, n = 9, 3 tanks for per diet with 3 fish randomly from each tank; i*n vitro*, n = 9, including three biological replicates and three technical replicates per biological replicate. The normality of data distribution and the homogeneity of variances were analyzed using the Kolmogorov-Smirnov test and Bartlett’s test, respectively. Differences between the two groups (Si-NC and Si-RNA group) were analyzed using an unpaired Student’s t-test, and multiple comparisons were assessed using a one-way analysis of variance (ANOVA) with Tukey’s multiple comparison test. Statistical significance was defined as a p-value < 0.05. Significance levels were denoted as follows: * *P*<0.05, ** *P*<0.01, *** *P*<0.001, and **** *P*<0.0001. Non-significant differences were labeled "ns".

## RESOURCE AVAILABILITY

### Data availability statement

The datasets in the current study are available from the corresponding author on reasonable request.

## Acknowledgments

This work was supported by the National Natural Science Foundation of China (grant No. 32273156; 32674059), Fundamental Research Funds for the Central Universities, China (grant No. 2662025SCPY003) and National Key Research and Development Program of China (grant No. 2025ZD0408100).

## Authors’ contributions

Song YF designed the experiment. Chi YJ conducted the experiment and data analysis with the help of Liu ZX, Feng GL and Hu NJ. Song YF drafted the manuscript. All the authors reviewed and approved the manuscript.

## Declarations

### Ethics approval and consent to participate

The study protocol was reviewed and approved by the Ethical Committee of Huazhong Agricultural University’s (HZAU) (identification code: Fish-2024-0127) and conformed to the ethical standards for the care and use of laboratory animals (no human subjects), as laid out in the 1964 Declaration of Helsinki and its later amendments.

## Supporting Information

### Supplementary Figures

**Figure. S1.**
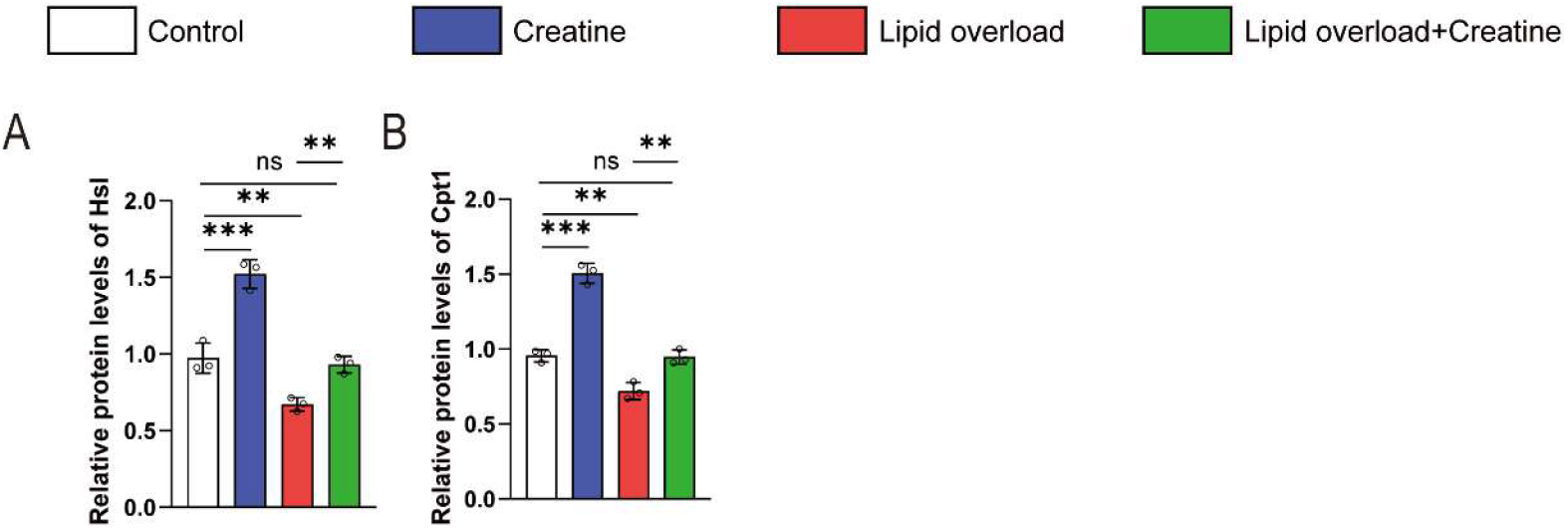
relative protein expression. (A) Quantification of HSL relative protein levels. (B) Quantification of Cpt1 relative protein levels.

**Figure. S2.**
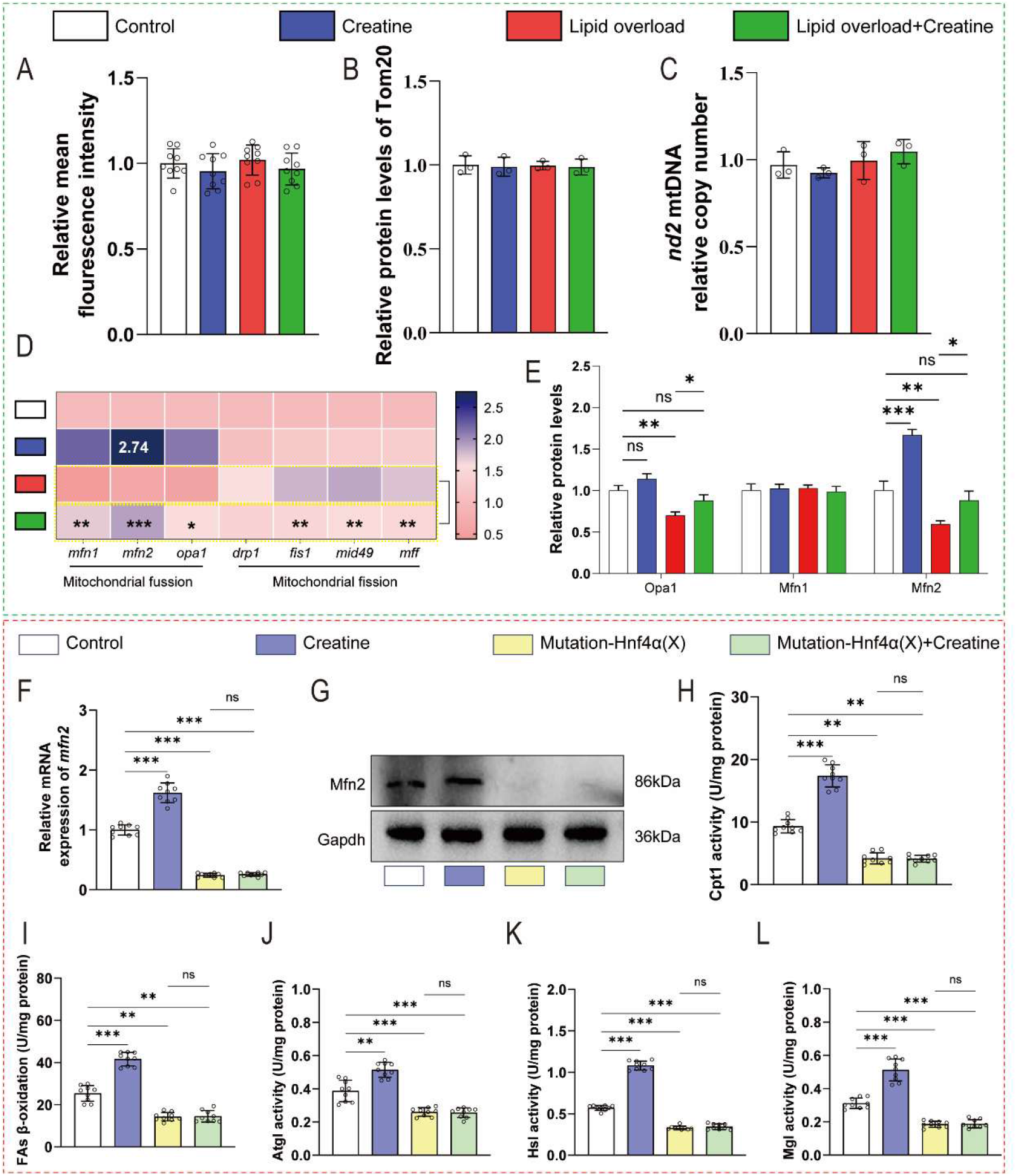
Dietary creatine promotes mitochondrial fusion and then increases PDM formation by activating the HNF4α-binding element in the mfn2 promoter. (A)Mitochondria content was quantified by flow cytometric analysis of FL3 (red) mean fluorescence intensity with MitoTracker Deep Red FM 644/665 staining. (B) Quantification of Tom20 relative protein levels. (C) Quantification of *nd2* mtDNA relative copy numbers. (D)The mRNA levels of genes involved in mitochondrial fusion and fission. (E) Quantification of Opa1, Mfn1 and Mfn2 relative protein levels. (F) The mRNA levels of *mfn2*. (G) Quantification of Mfn2 relative protein levels. (H) Cpt1 activity of isolated mitochondria. (I) FAs *β*-oxidation rate of isolated mitochondria. (J) Atgl activity. (K) HSL activity. (L) Mgl activity.

**Figure. S3.**
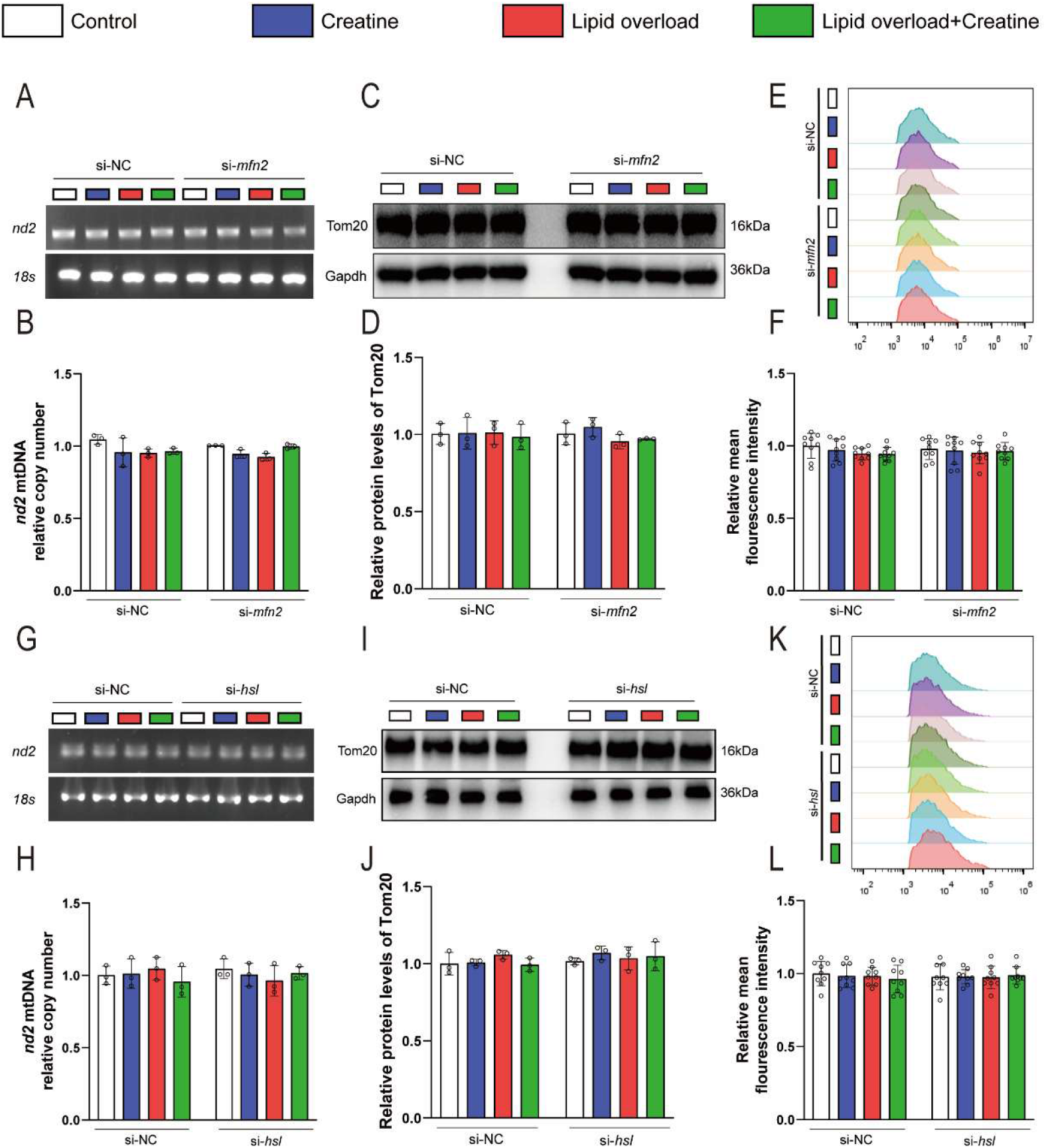
si-*mfn2* and si-*hsl* did not significantly alter mitochondrial number. (A) mtDNA fold-change analysis in primary hepatocytes treated with si-NC or si-*mfn2* for 48 h (*in vitro* experiments). (B)Quantification of *nd2* mtDNA relative copy number in panel A. (C)Western blot analysis of Tom20 in primary hepatocytes treated with si-NC or si-*mfn2* for 48 h (*in vitro* experiments). (D) Quantification of Tom20 relative protein levels in panel C. (E) Flow cytometry analysis of mitochondrial content in primary hepatocytes treated with si-NC or si-*mfn2* for 48 h (*in vitro* experiments). (F) Mitochondria content was quantified by flow cytometric analysis of FL3 (red) mean fluorescence intensity with MitoTracker Deep Red FM 644/665 staining. (G) mtDNA fold-change analysis in primary hepatocytes treated with si-NC or si-*hsl* for 48 h (*in vitro* experiments). (H)Quantification of *nd2* mtDNA relative copy number in panel A. (I)Western blot analysis of Tom20 in primary hepatocytes treated with si-NC or si-*hsl* for 48 h (*in vitro* experiments). (J) Quantification of Tom20 relative protein levels in panel C. (K) Flow cytometry analysis of mitochondrial content in primary hepatocytes treated with si-NC or si-*hsl* for 48 h (*in vitro* experiments). (L) Mitochondria content was quantified by flow cytometric analysis of FL3 (red) mean fluorescence intensity with MitoTracker Deep Red FM 644/665 staining.

**Figure. S4.**
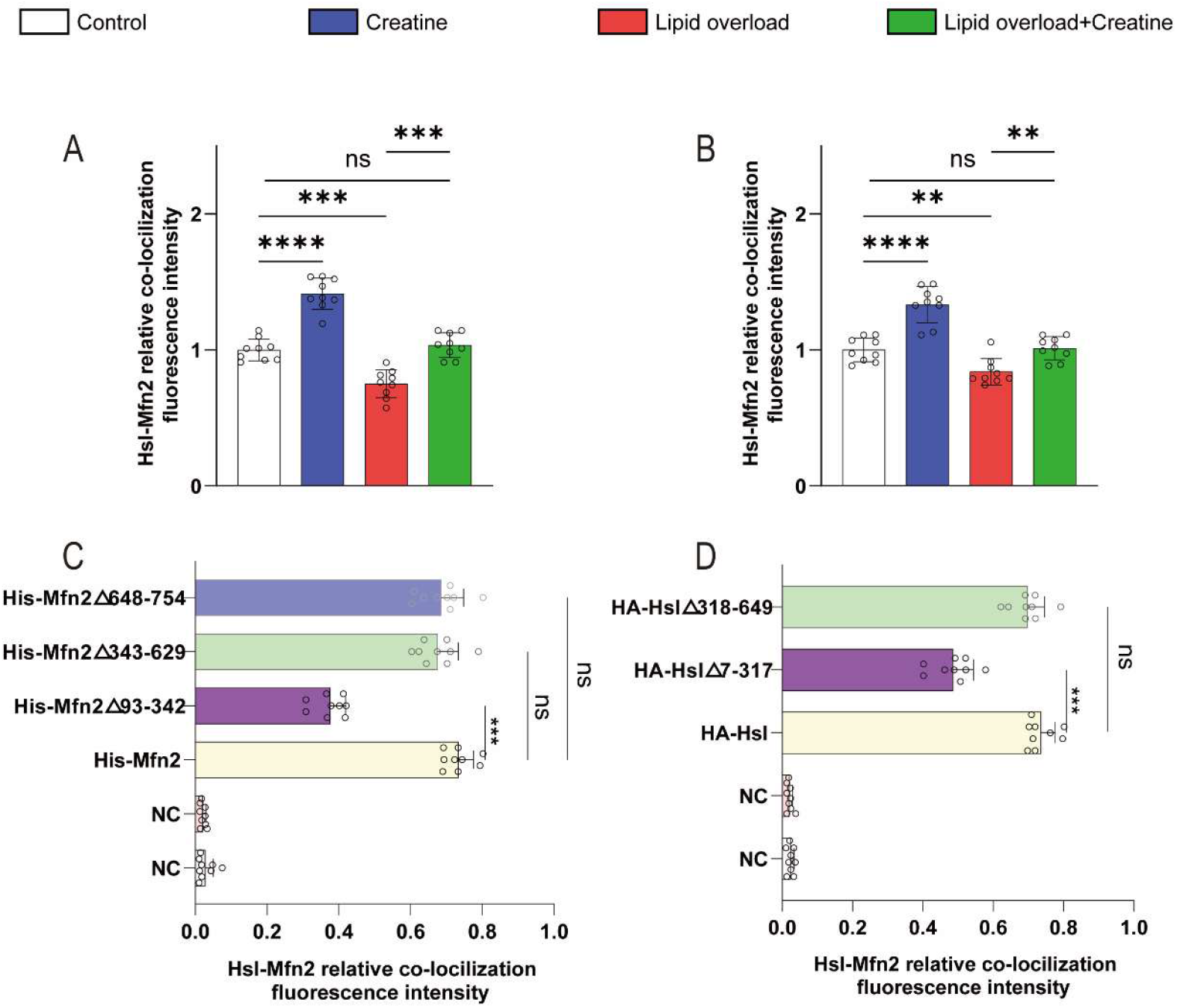
HSL-Mfn2 relative co-locilization fluorerscence intensity. (A) Quantification of the relative colocalization fluorescence intensity of HSL-Mfn2 in liver tissues. (B) Quantification of the relative colocalization fluorescence intensity of HSL-Mfn2 in primary hepatocytes. (C) Quantification of the relative colocalization fluorescence intensity of HSL-Mfn2 in 293T cells transfected with wild-type Mfn2 and their truncated mutants. (D) Quantification of the relative colocalization fluorescence intensity of HSL-Mfn2 in 293T cells transfected with wild-type HSL and their truncated mutants.

**Figure. S5.**
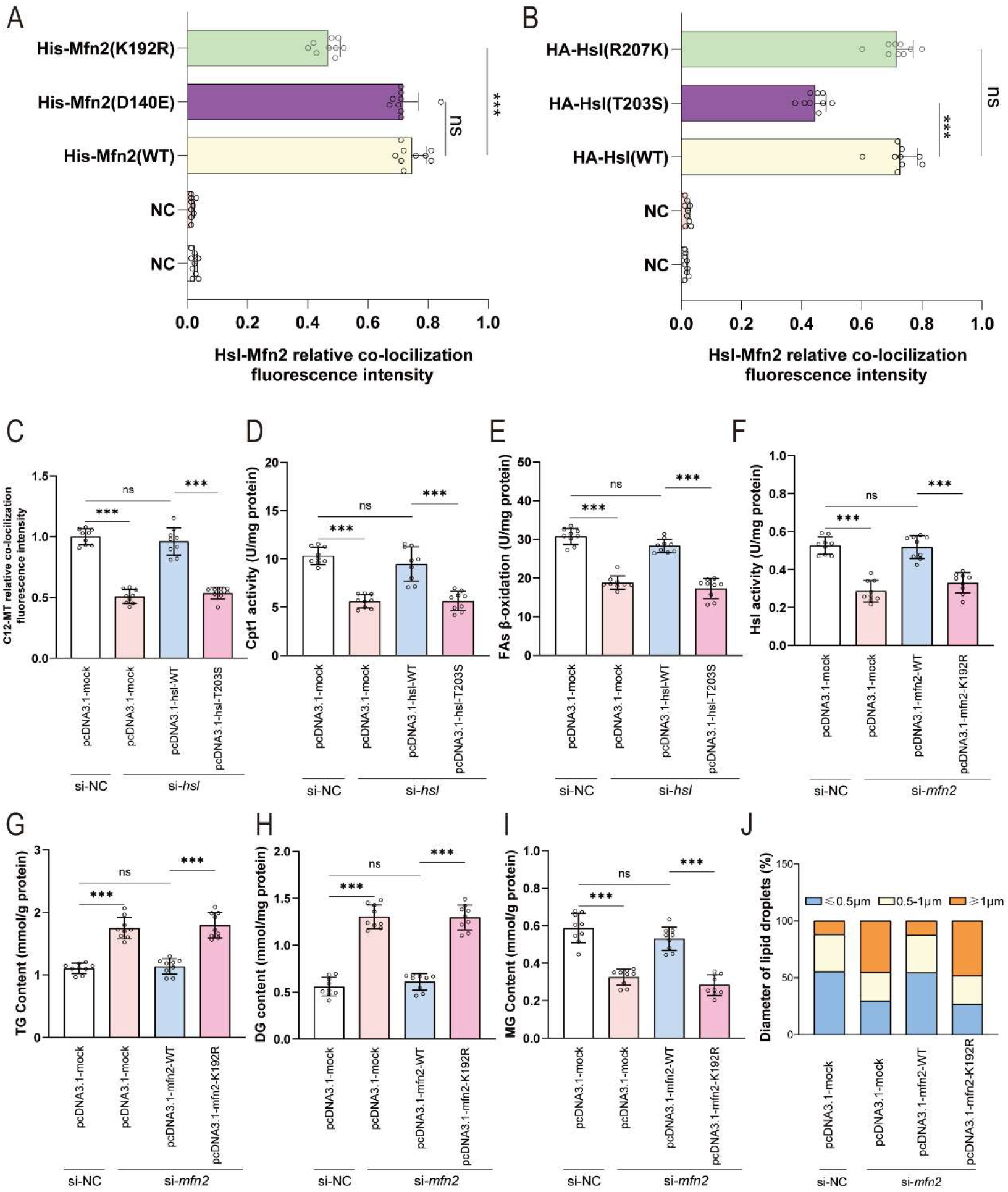
Regulatory function of Mfn2-HSL-governed mitochondria-lipid droplet coupling in lipid droplet lipolysis. (A) Quantification of the relative colocalization fluorescence intensity of HSL-Mfn2 in 293T cells transfected with wild-type Mfn2 and their truncated mutants. (B) Quantification of the relative colocalization fluorescence intensity of HSL-Mfn2 in 293T cells transfected with wild-type HSL and their truncated mutants. (C) The fusion rate between mitochondria and Red C12 in panel H. (D) CPT1 activity in mitochondria isolated from 293T cells transfected with wild-type *hsl* and its mutants. (E) FAs *β*-oxidation rate in mitochondria isolated from 293T cells transfected with wild-type *hsl* and its mutants. (F) HSL activity in 293T cells transfected with wild-type *mfn2* and its mutants. (G) TG in 293T cells transfected with wild-type *mfn2* and its mutants. (H) DG in 293T cells transfected with wild-type *mfn2* and its mutants. (I) MG in 293T cells transfected with wild-type *mfn2* and its mutants. (J) Quantification of lipid droplet diameter in panel I.

**Figure. S6.**
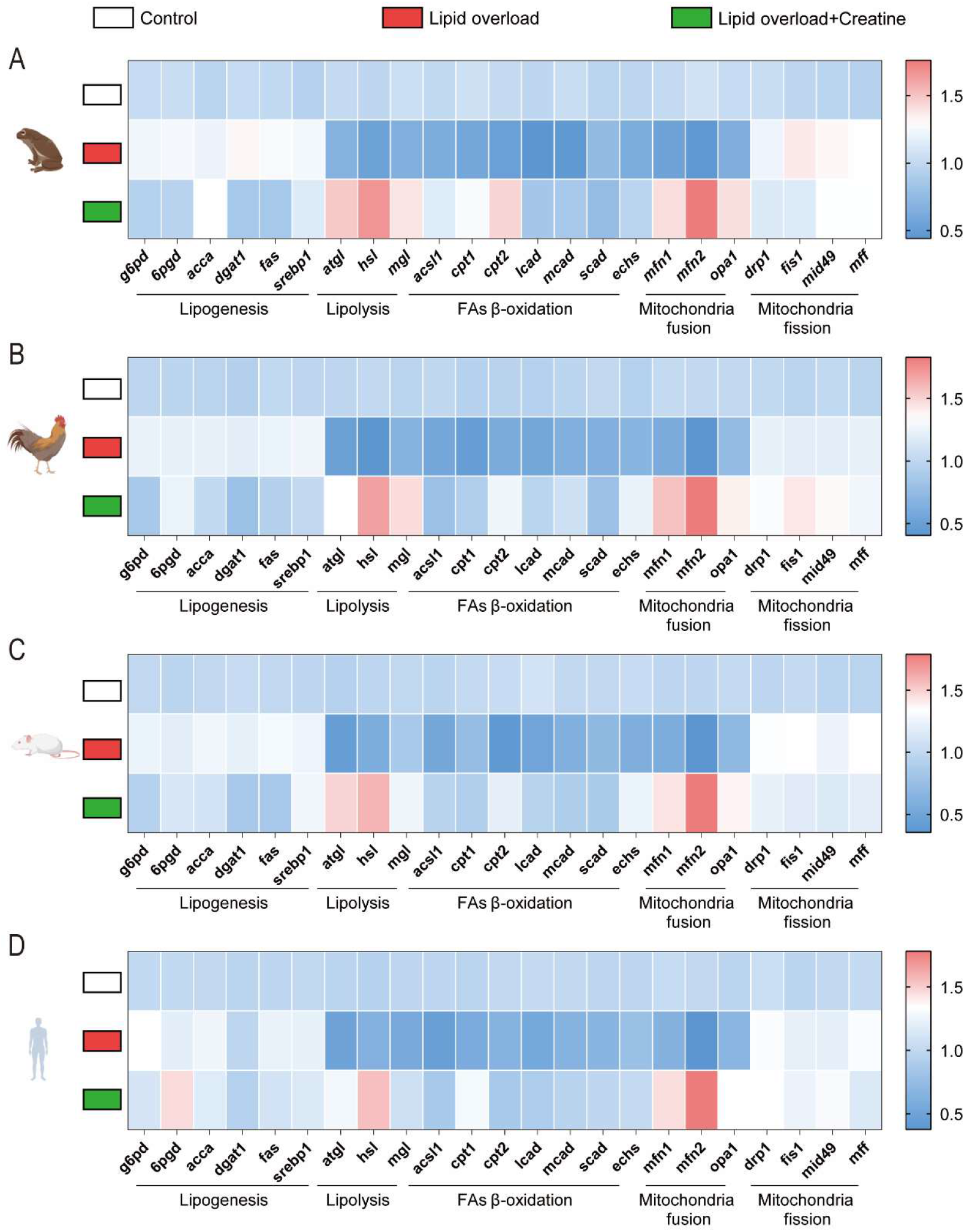
The mRNA levels expression from frog to human. A) The mRNA levels of genes involved in lipid metabolism and mitochondrial fusion and fission in hepatocytes from frog. B) The mRNA levels of genes involved in lipid metabolism and mitochondrial fusion and fission in hepatocytes from chicken. C) The mRNA levels of genes involved in lipid metabolism and mitochondrial fusion and fission in hepatocytes from mice. D) The mRNA levels of genes involved in lipid metabolism and mitochondrial fusion and fission in hepatocytes from human. For all data, in vitro, n = 9, including three biological replicates and three technical replicates per biological replicate. Statistical significance was defined as a p-value < 0.05. Significance levels were denoted as follows: * P<0.05, ** P<0.01, *** P<0.001, and **** P<0.0001. Non-significant differences were labeled "ns".

**Figure. S7.**
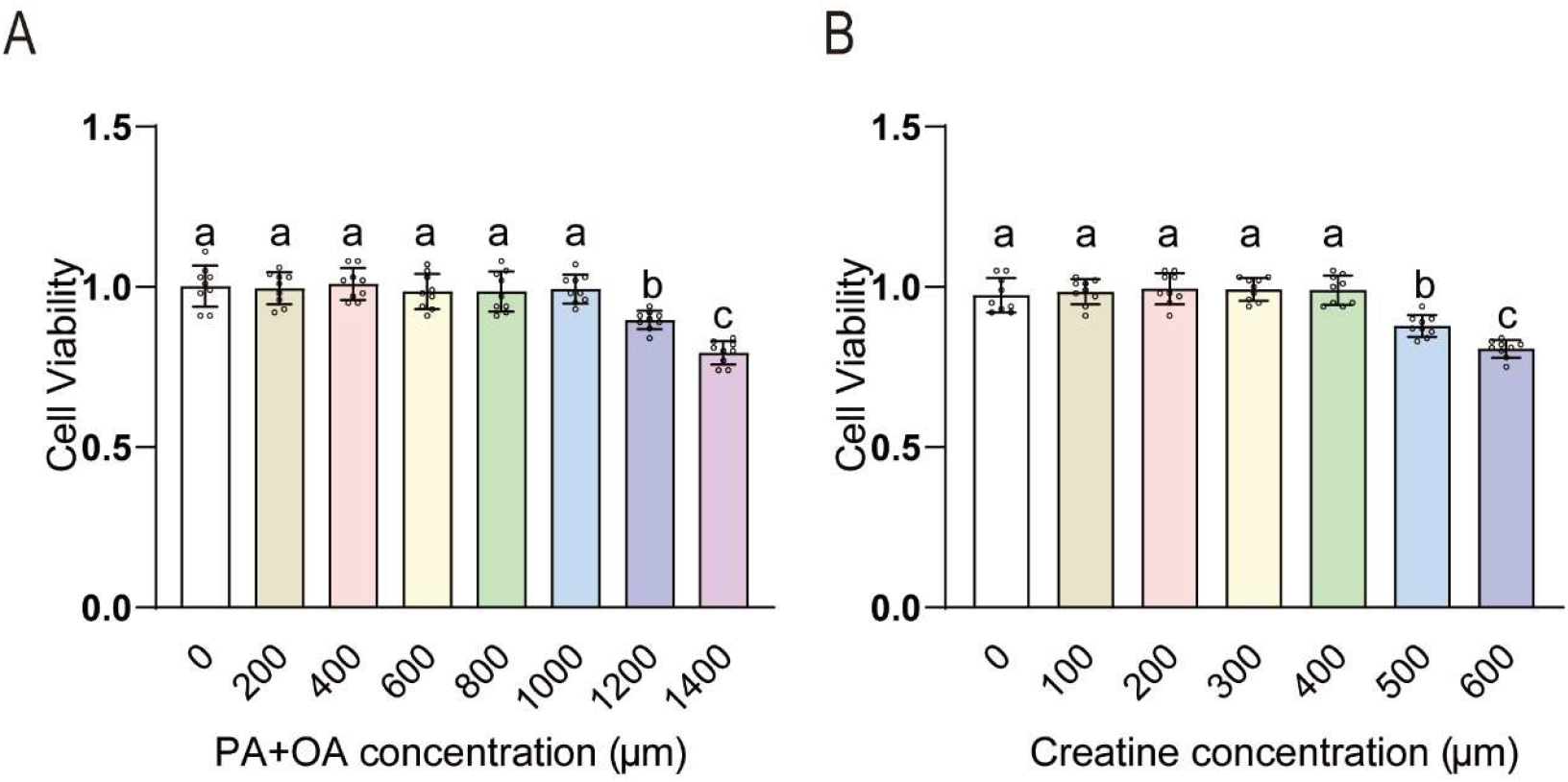
Cell viability in primary hepatocytes isolated from yellow catfish. A, B) Cell viability of hepatocytes incubated with OA:PA/1:1 or Creatine 48 h using the MTT assay. All data were expressed as mean ± S.E.M. n = 9, *in vitro* each treatment was performed in triplicate plates and then at least three biological replicates (3-well cells) for one plate, *P <* 0.05. Values without the same letter indicate significant differences among different treatments.

**Figure. S8.**
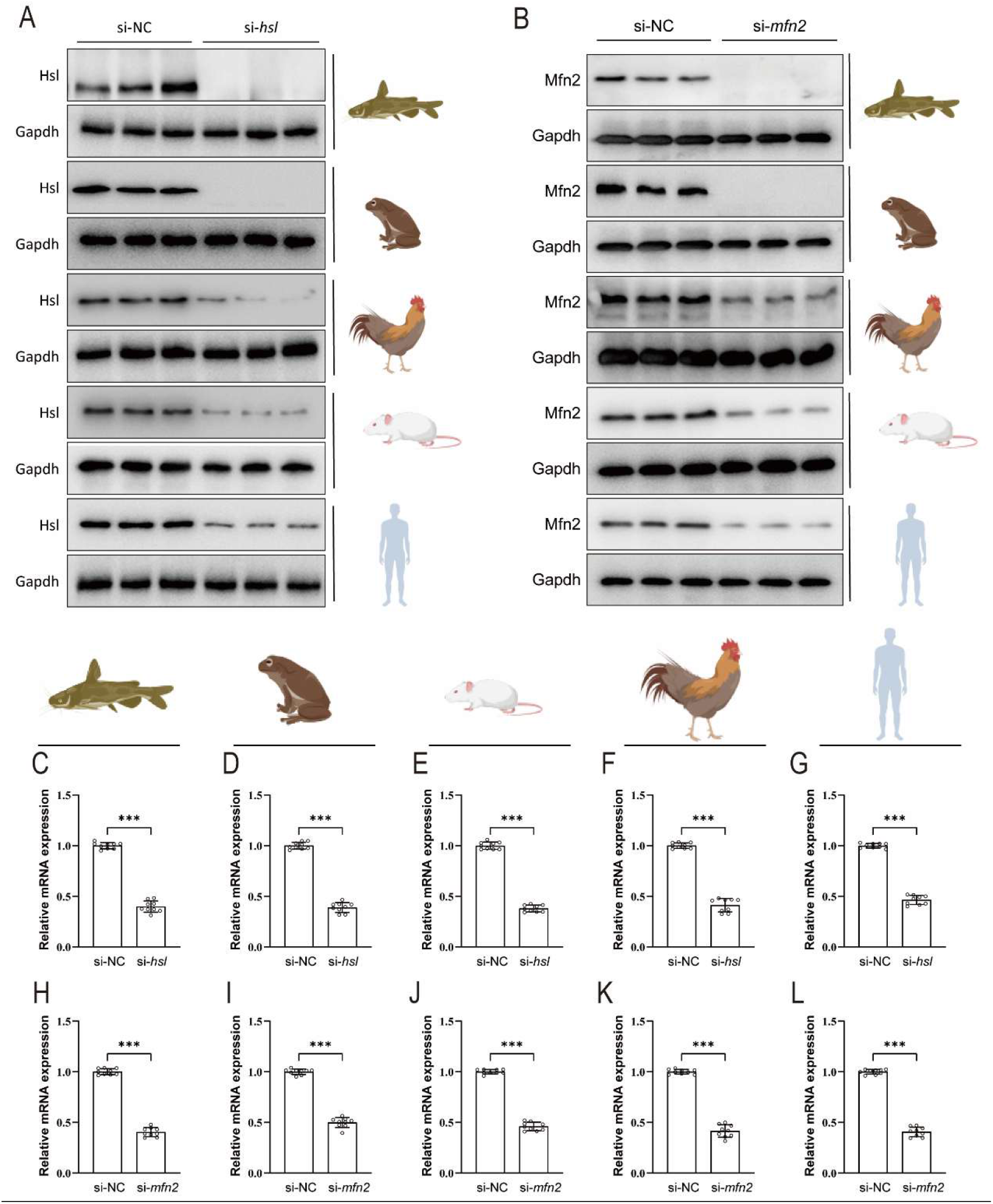
siRNA interference efficiency in primary hepatocytes (isolated from yellow catfish and frog) and hepatocyte lines (from chicken, mouse and human). A)The protein levels of HSL in primary hepatocytes treated with si-NC or si-mfn2 for 48h. B)The protein levels of Mfn2 in primary hepatocytes treated with si-NC or si-mfn2 for 48h. C-G) The mRNA levels of *hsl* in primary hepatocytes treated with si-NC or si-mfn2 for 48h. H-L) The mRNA levels of *mfn2* in primary hepatocytes treated with si-NC or si-mfn2 for 48h. All data were expressed as mean ± S.E.M. For western blot, n=3 three biological replicates for one treatment. For other data, n = 9, *in vitro* each treatment was performed in triplicate plates and then at least three biological replicates (3-well cells) for one plate. Significance levels were denoted as ∗P < 0.05.

### Supplementary tables

**Table S1.** Growth performance, morphological parameters and feed utilization.

|  | Control | Creatine | Lipid overload | Lipid overload<br>+ Creatine |
| --- | --- | --- | --- | --- |
| Survival, % | 100±0.00 | 100±0.00 | 100±0.00 | 100±0.00 |
| IBW, g/fish | 5.20±0.02 | 5.21±0.01 | 5.21±0.01 | 5.24±0.01 |
| FBW, g/fish | 21.48±0.08<sup>b</sup> | 22.46±0.13<sup>c</sup> | 19.93±0.10<sup>a</sup> | 21.72±0.14<sup>b</sup> |
| WG, % | 313.19±0.32<sup>b</sup> | 331.44±0.54<sup>c</sup> | 282.95±0.25<sup>a</sup> | 314.28±0.37<sup>b</sup> |
| SGR, %/d | 2.53±0.01<sup>b</sup> | 2.61±0.01<sup>c</sup> | 2.40±0.01<sup>a</sup> | 2.54±0.02<sup>b</sup> |
| FCR | 1.25±0.01<sup>c</sup> | 1.14±0.01<sup>a</sup> | 1.35±0.02<sup>d</sup> | 1.21±0.01<sup>b</sup> |

<sup>a)</sup>All data were expressed as mean ± S.E.M. (n=3), *P* < 0.05. Values without the same letter indicate significant difference among different treatments.
IBW (g/fish), initial mean body weight;
FBW (g/fish), final mean body weight;
WG (weight gain, %) = (FBW-IBW)/IBW×100;
SGR (specific growth rate, % /d) = 100× [ln (FBW)-ln (IBW)]/d;
FCR (feed conversion rate) = dry food fed (g)/wet weight gain (g).

**Table S2.**
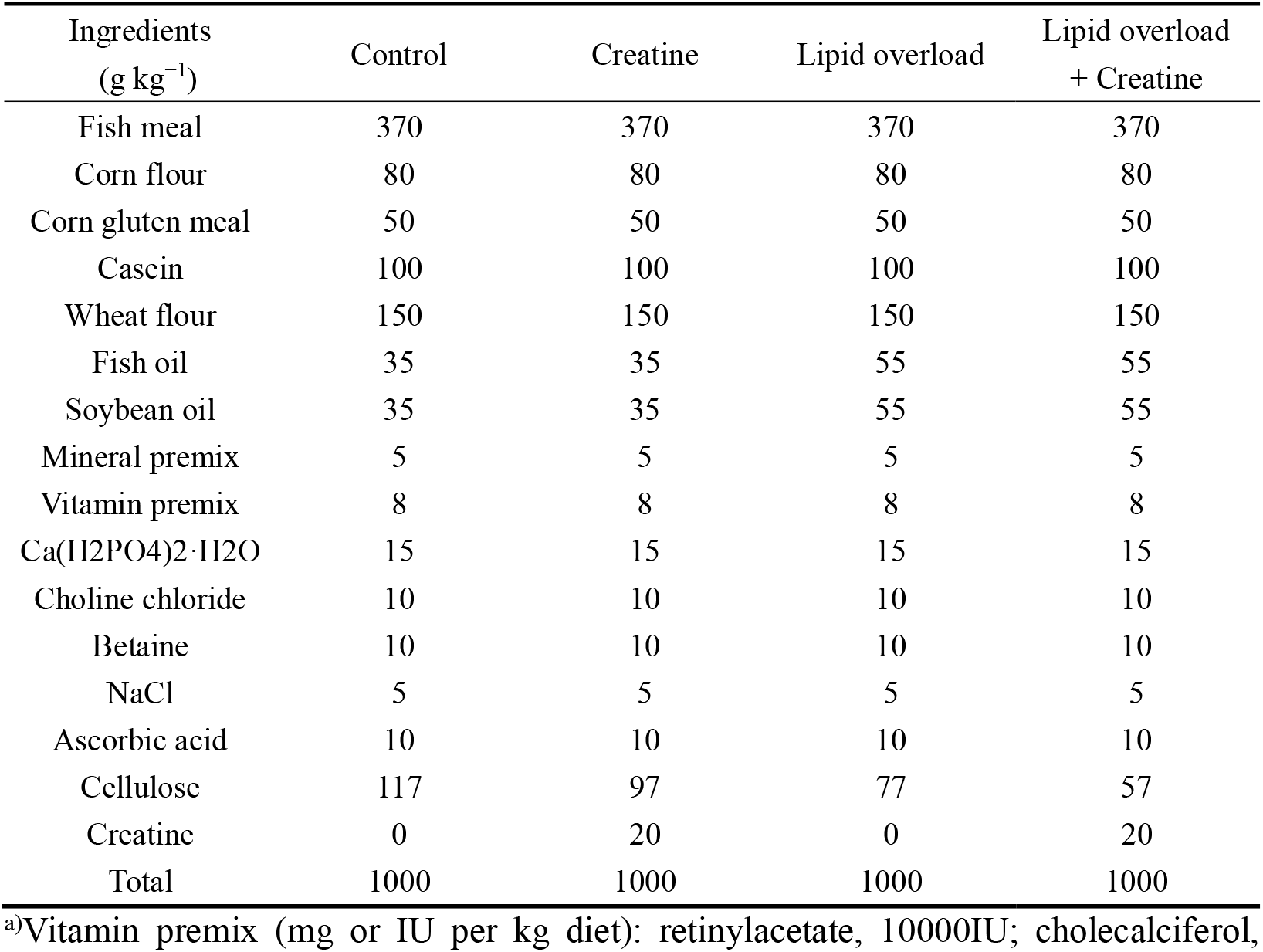
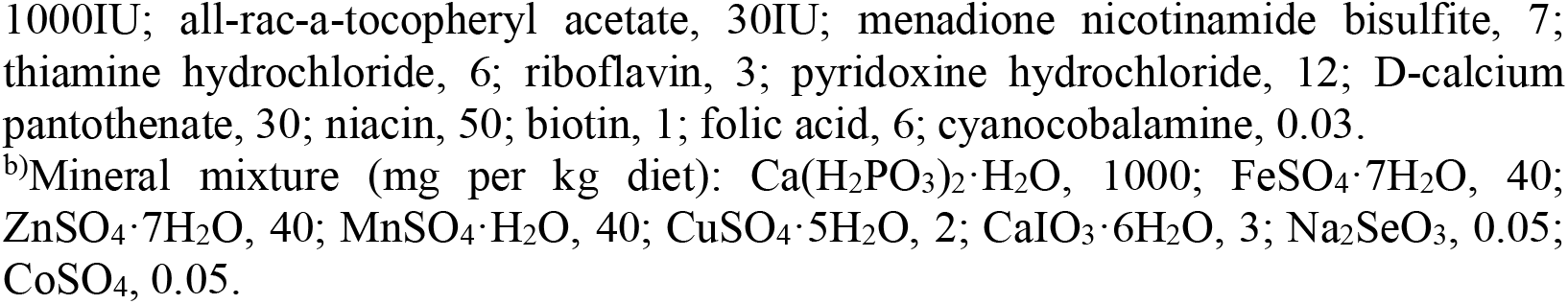
Ingredients of experimental diets.

**Table S3.**
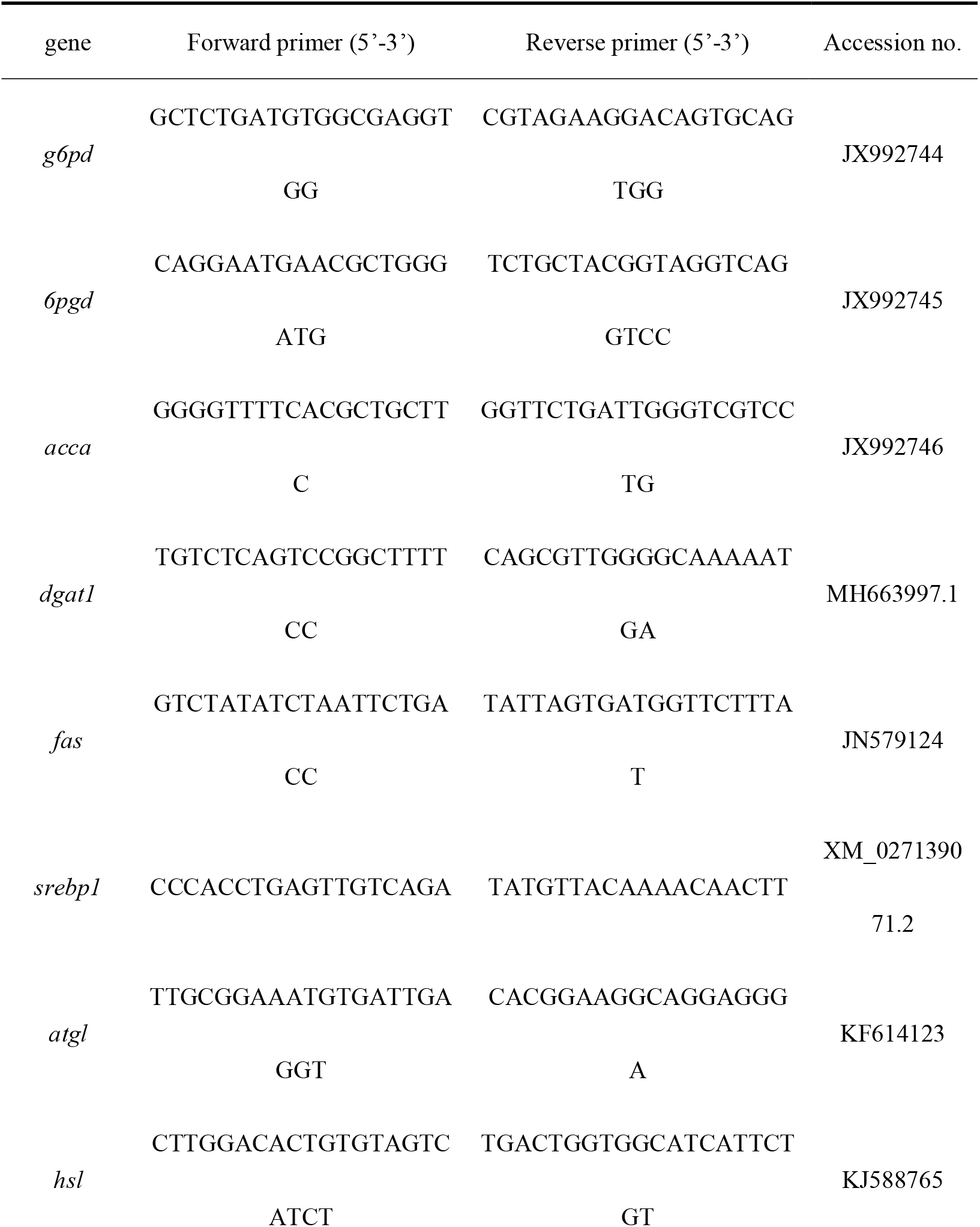
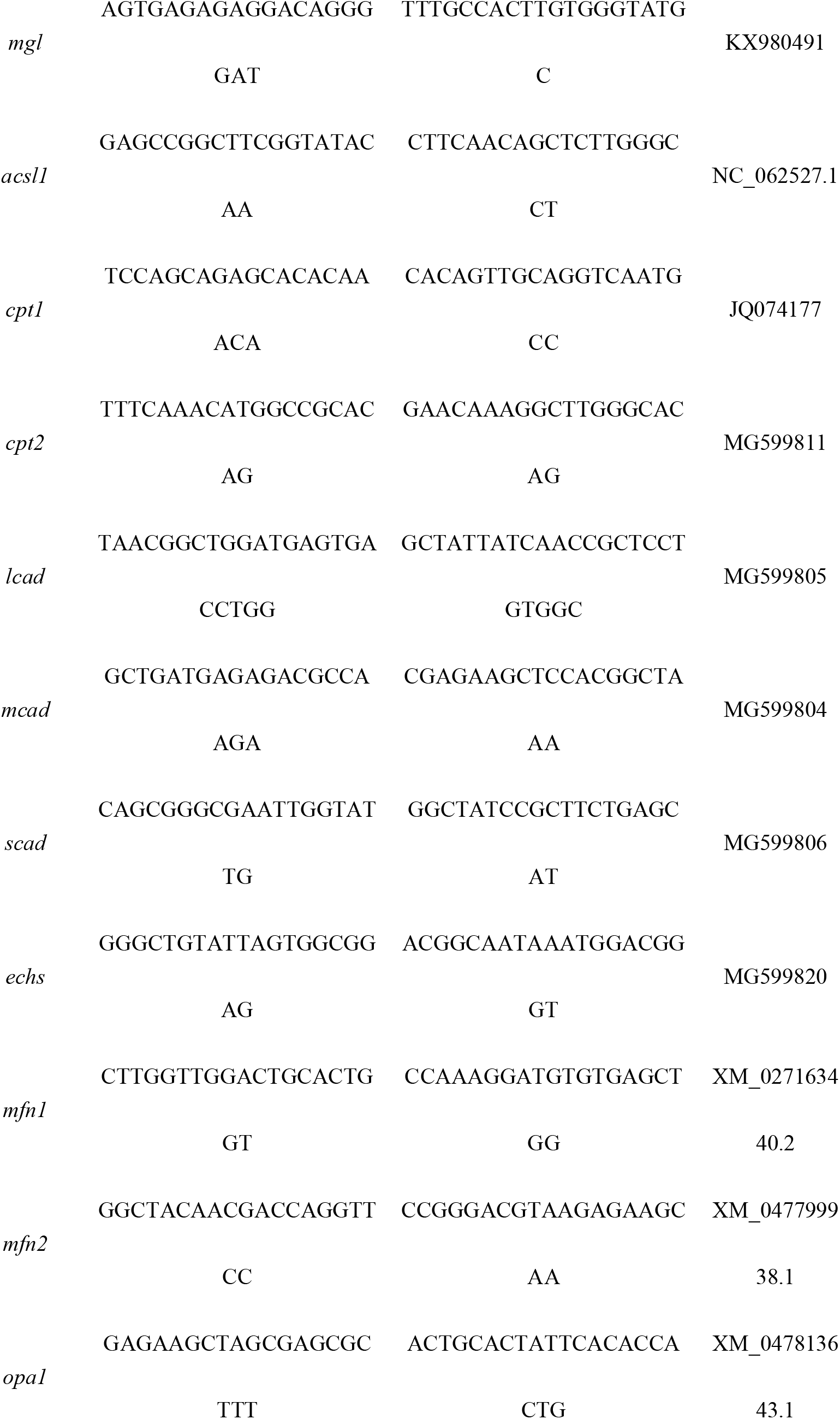
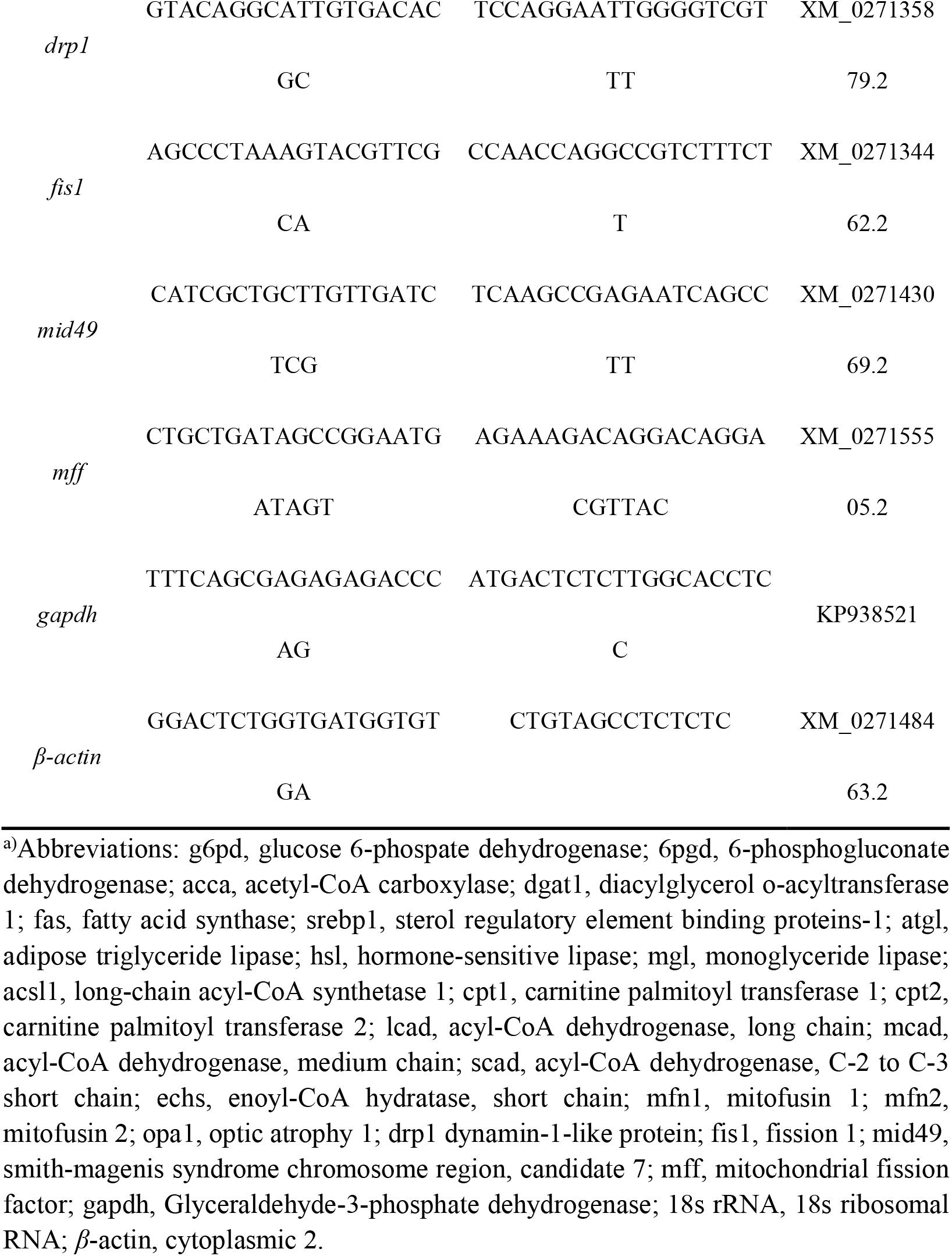
Primers in yellow catfish for quantitative real-time PCR analysis.

**Table S4.**
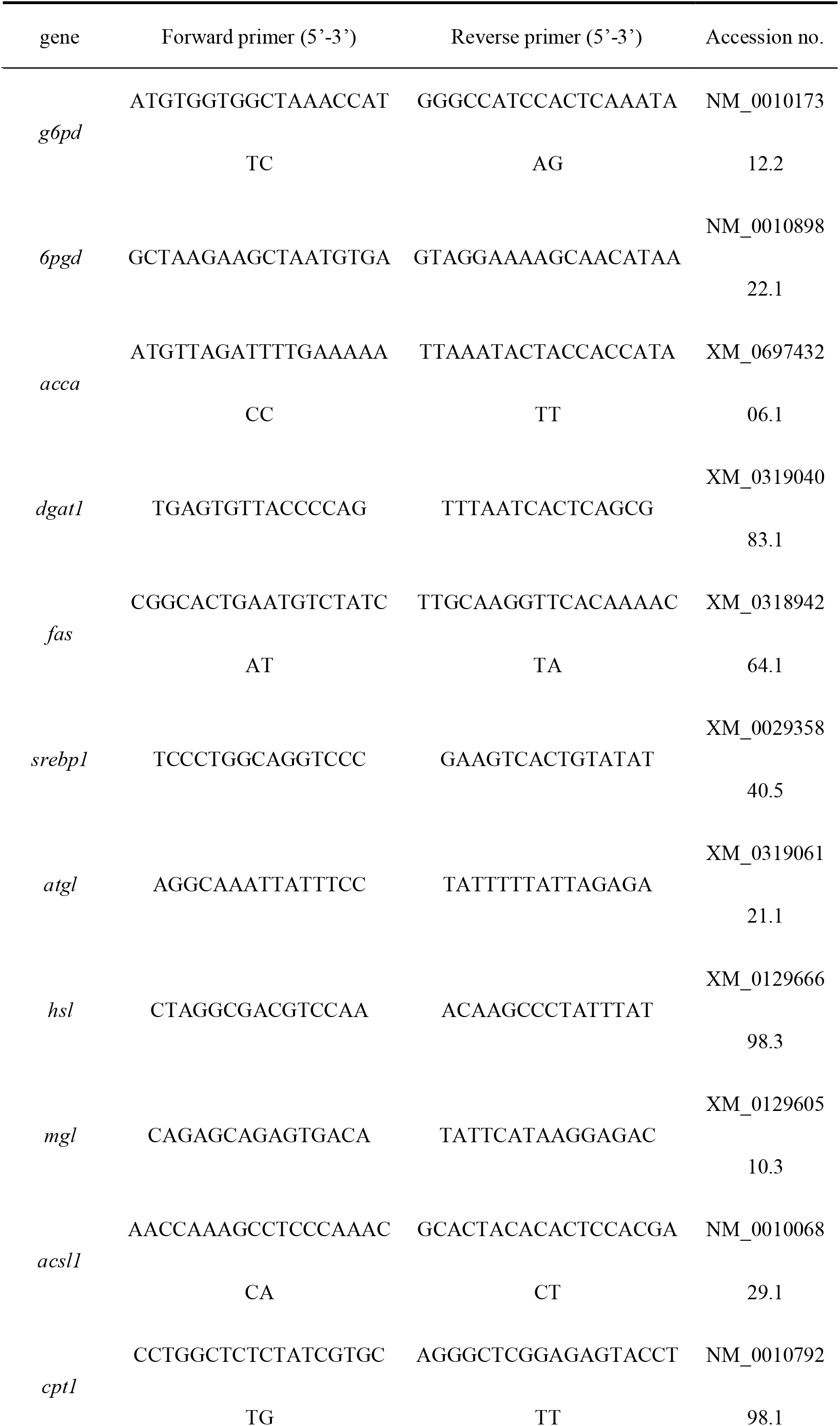
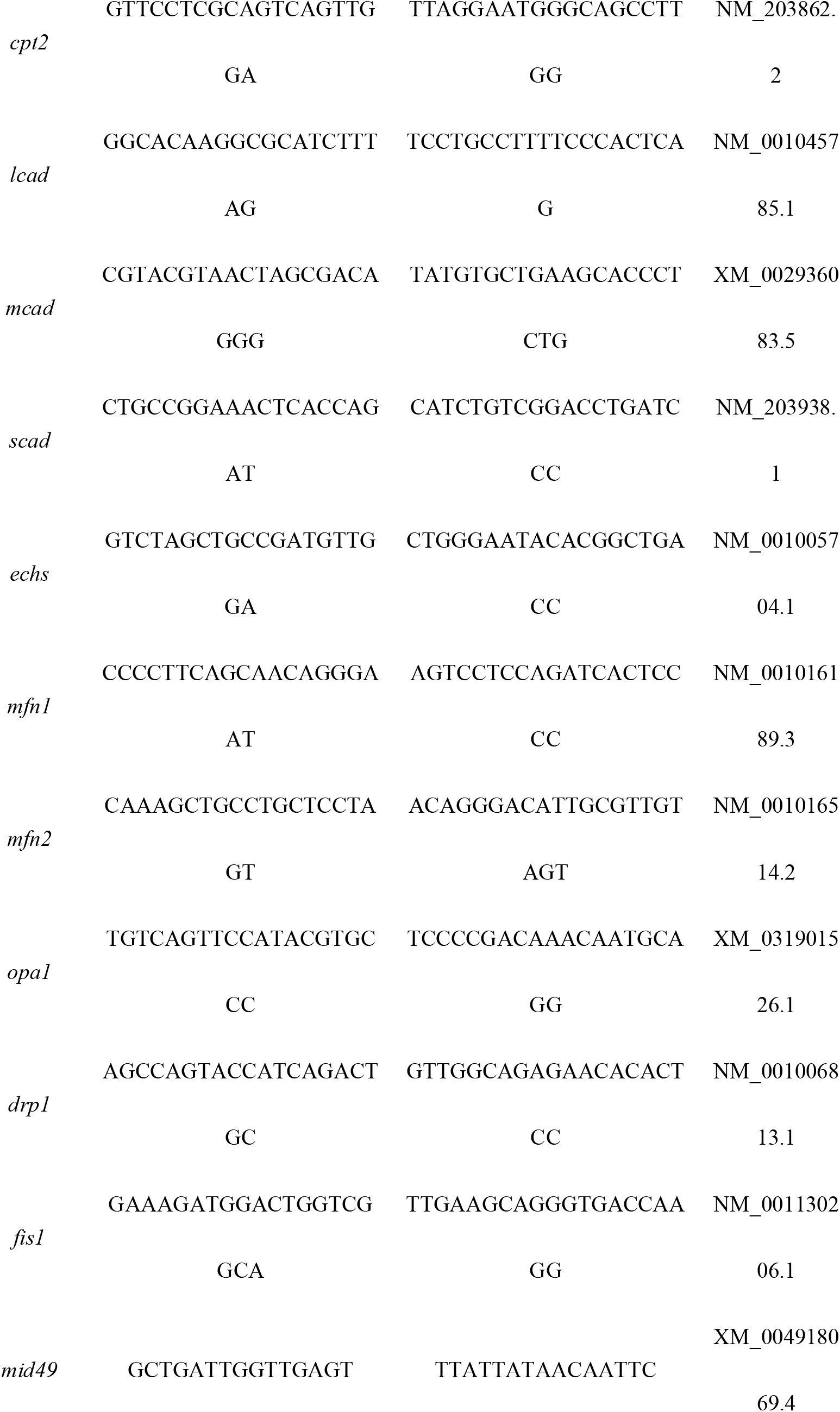
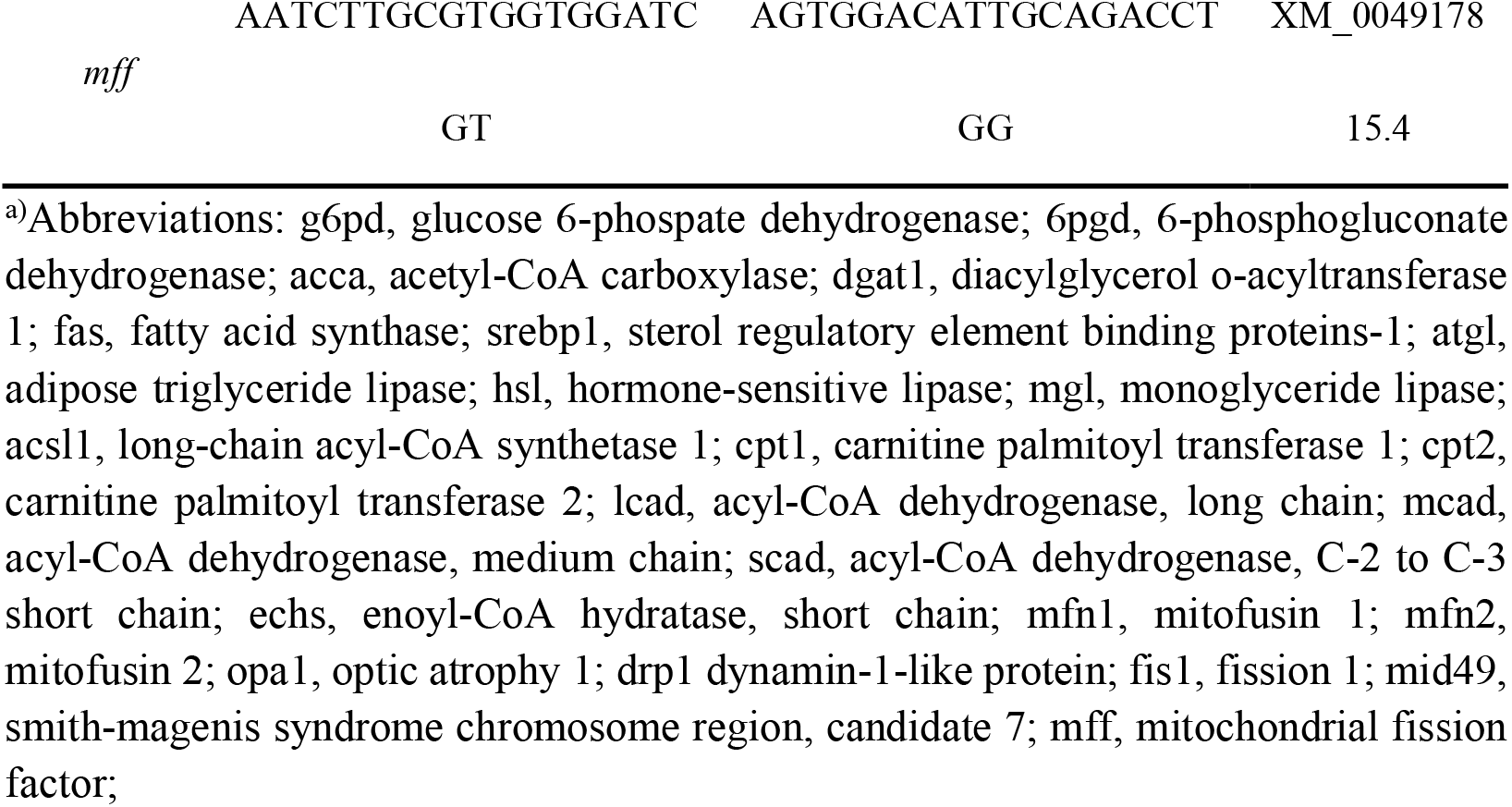
Primers in frog for quantitative real-time PCR analysis.

**Table S5.**
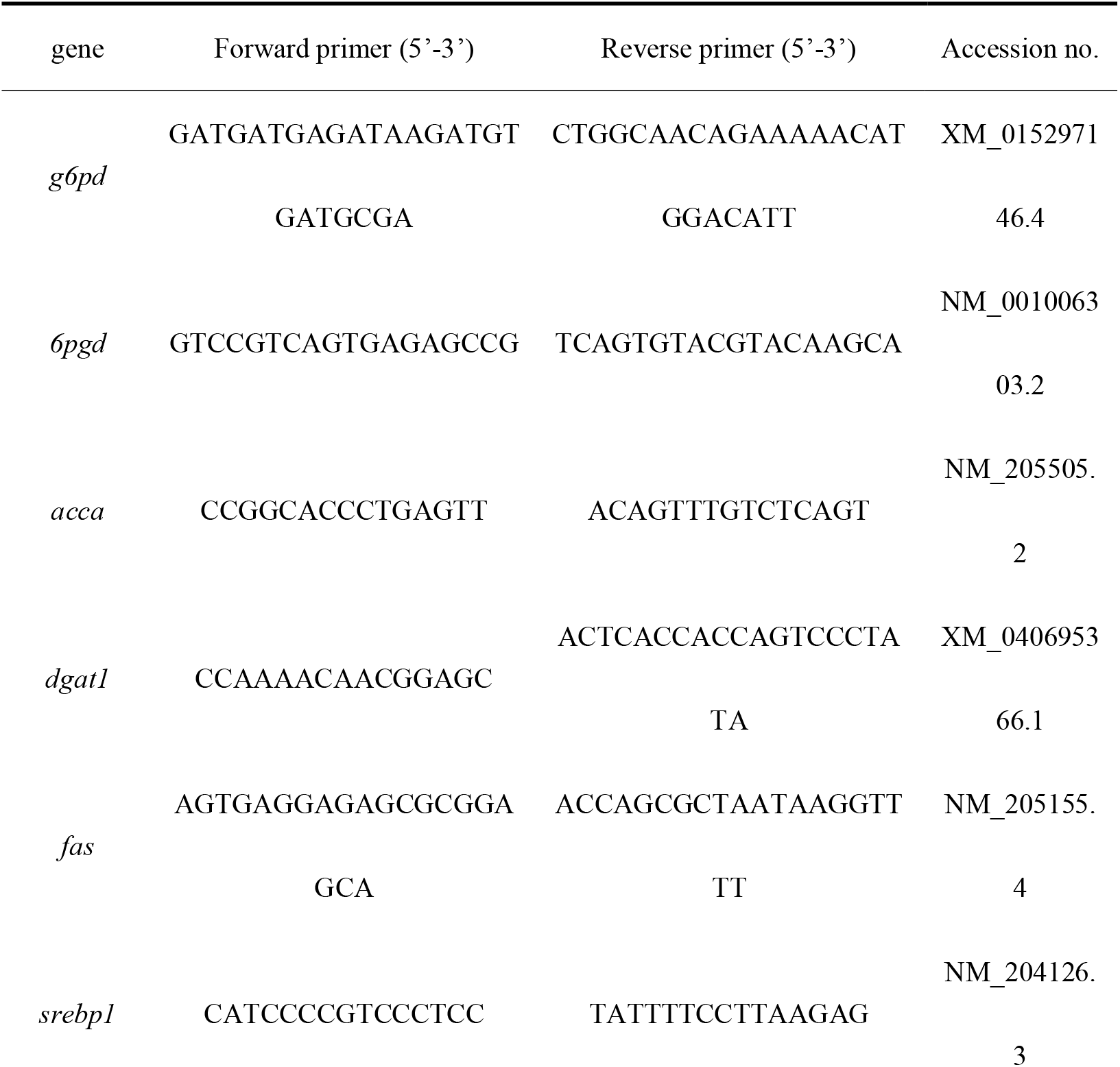
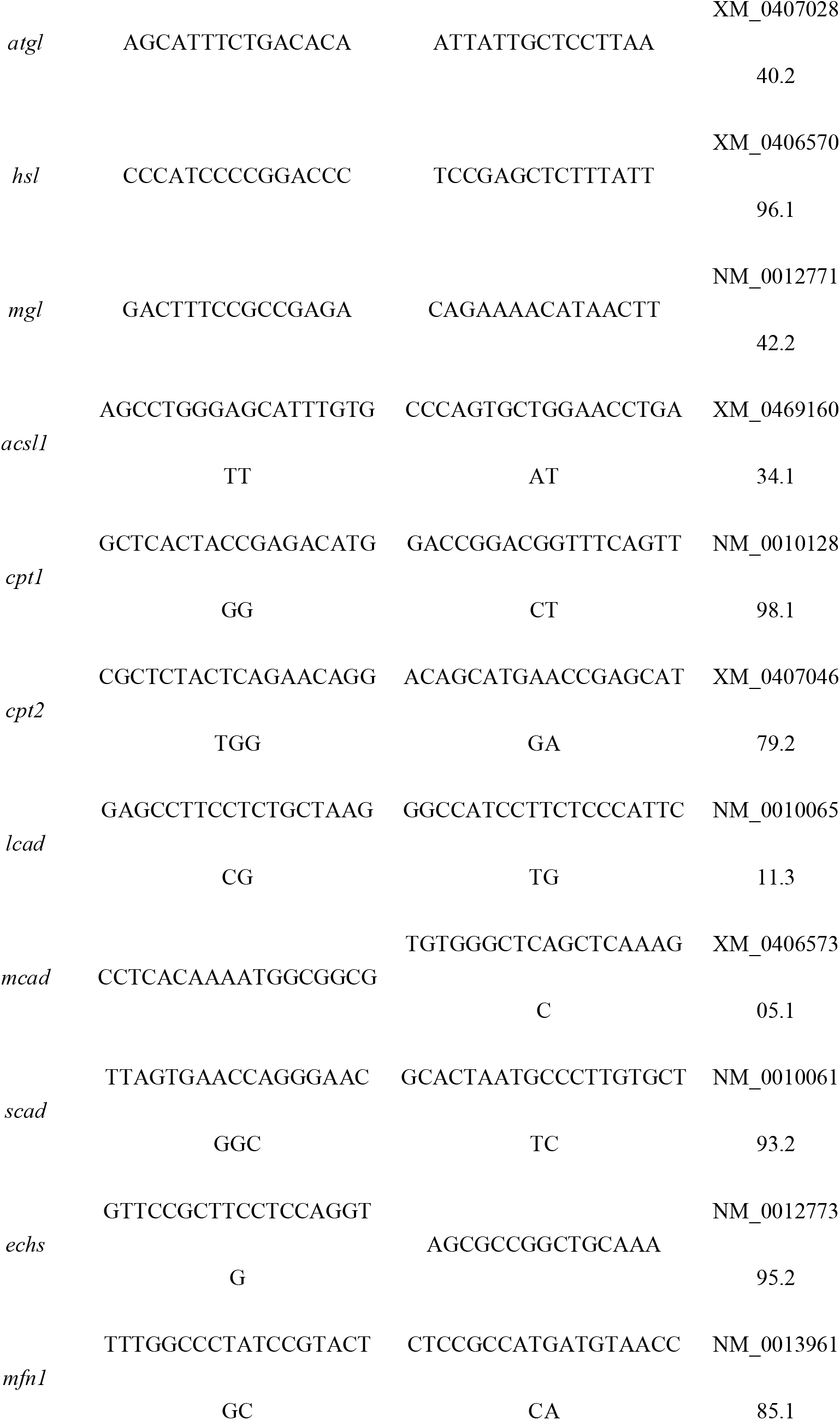
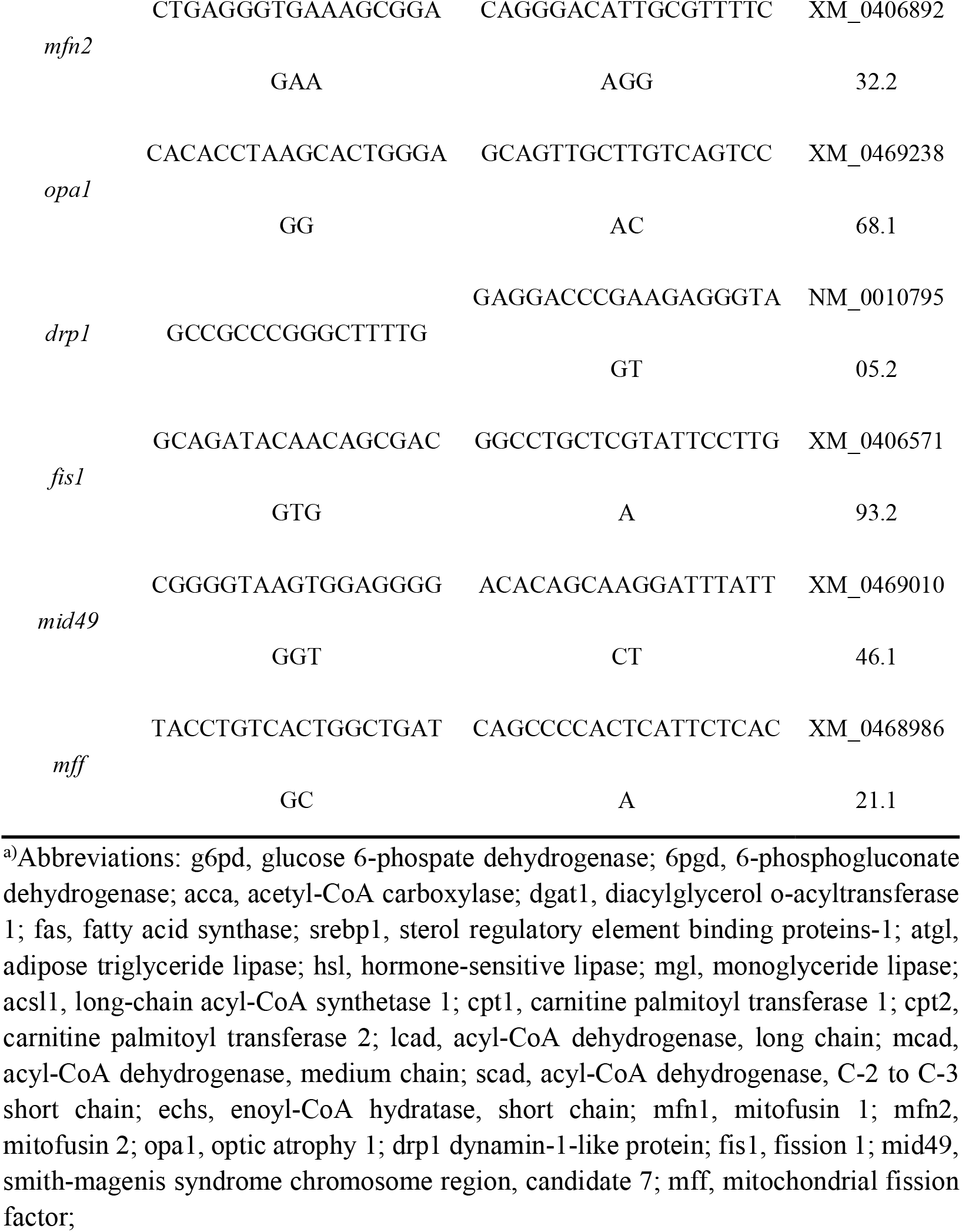
Primers in chicken for quantitative real-time PCR analysis.

**Table S6.**
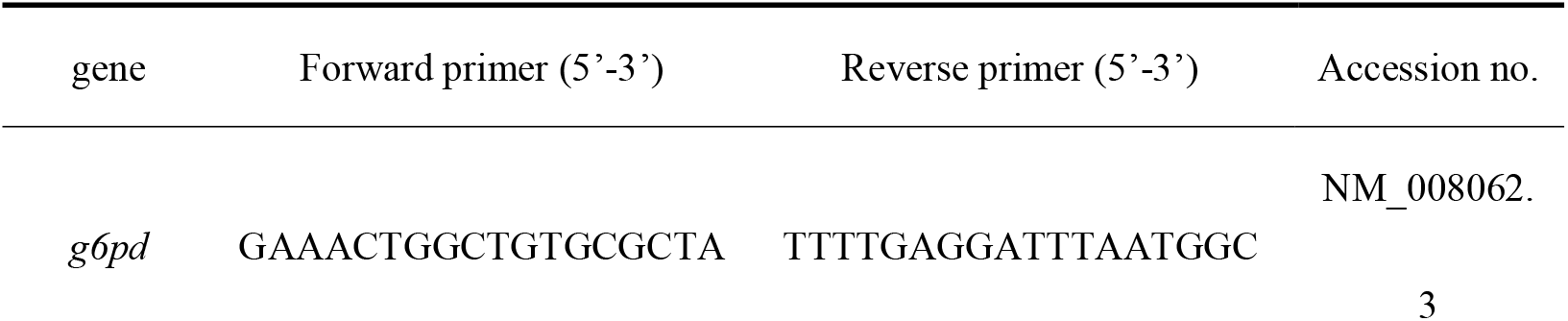
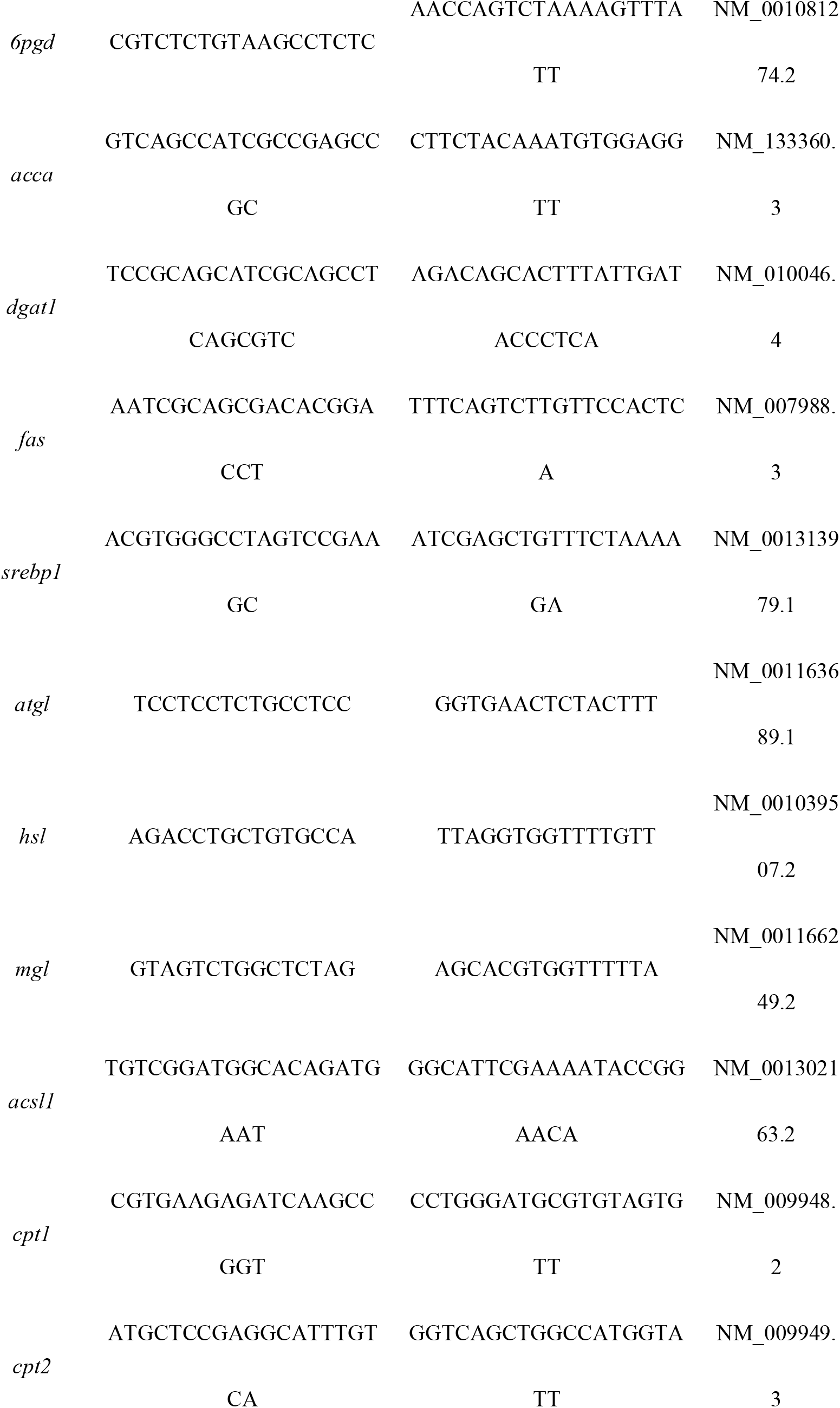
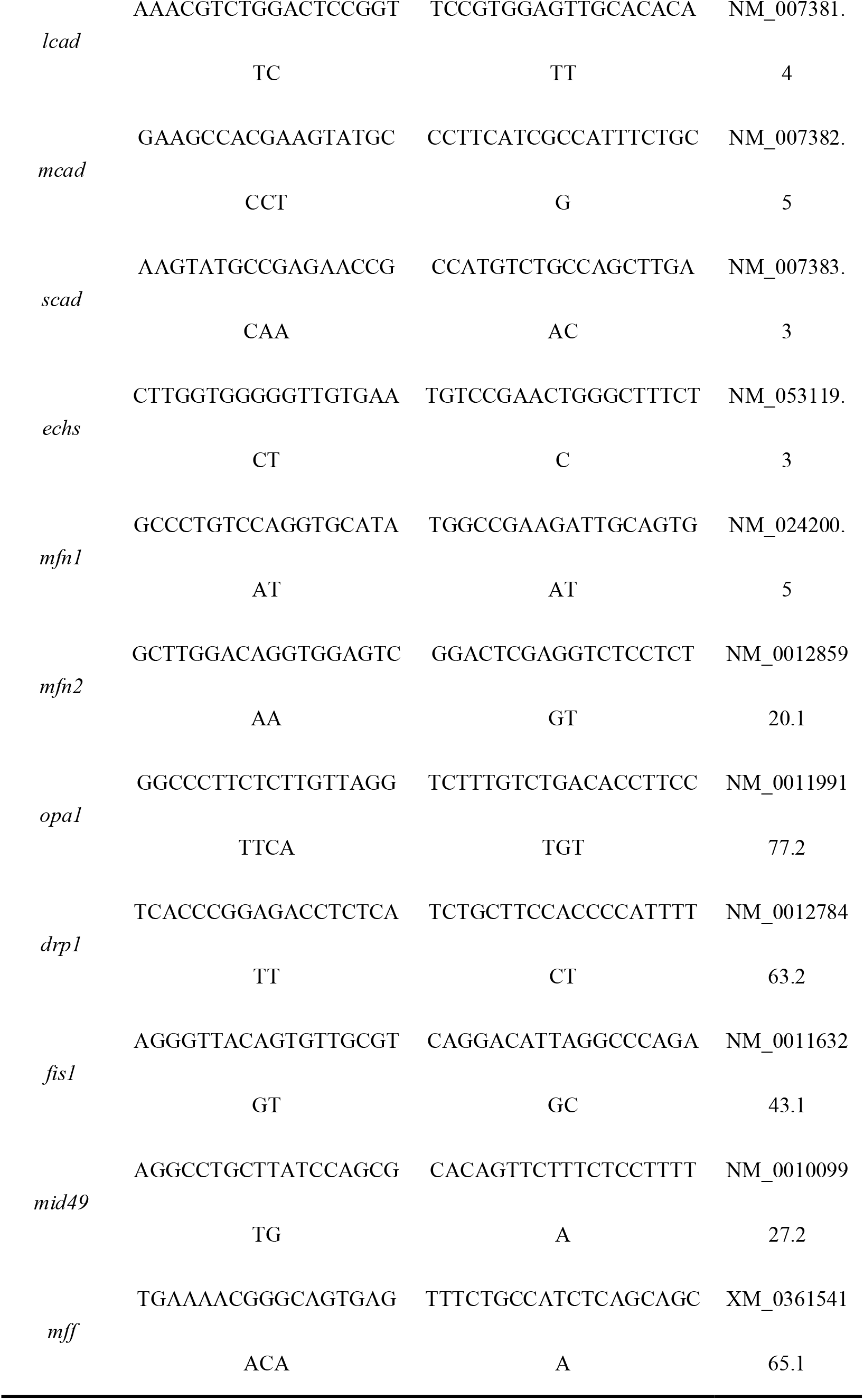
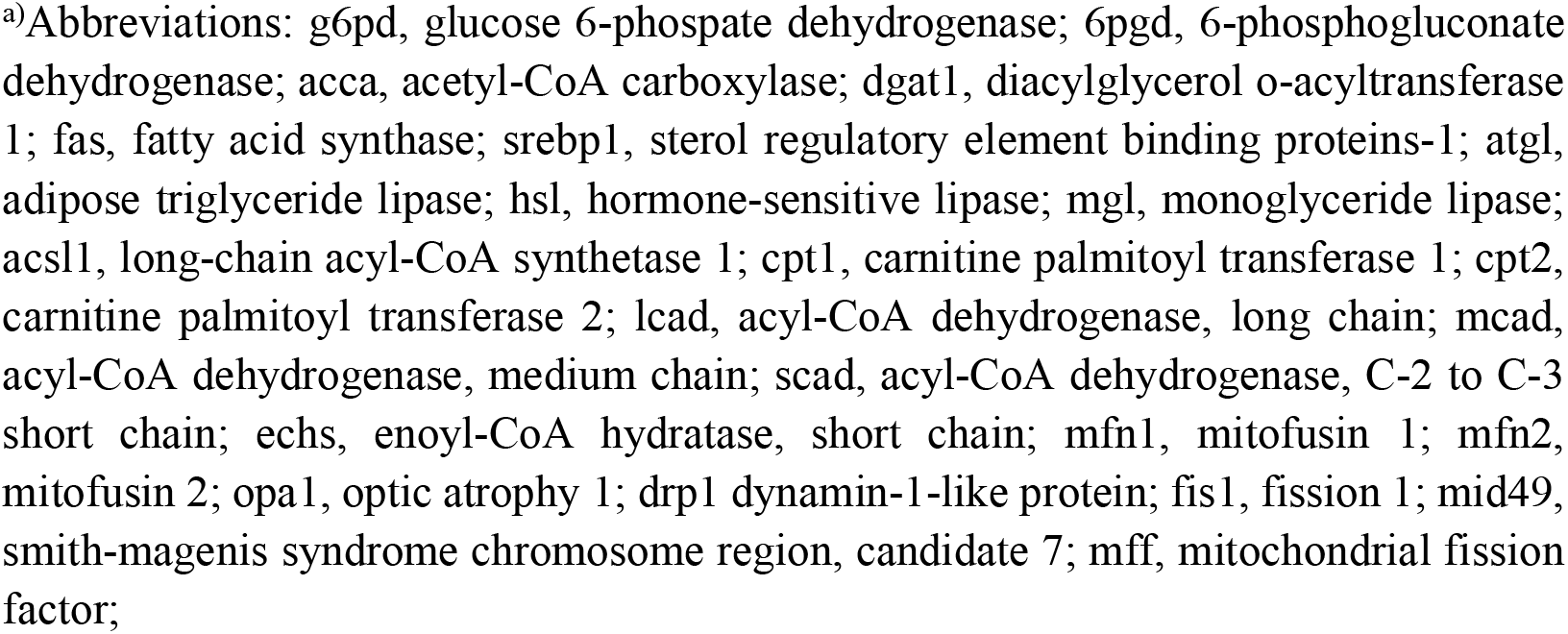
Primers in mouse for quantitative real-time PCR analysis.

**Table S7.**
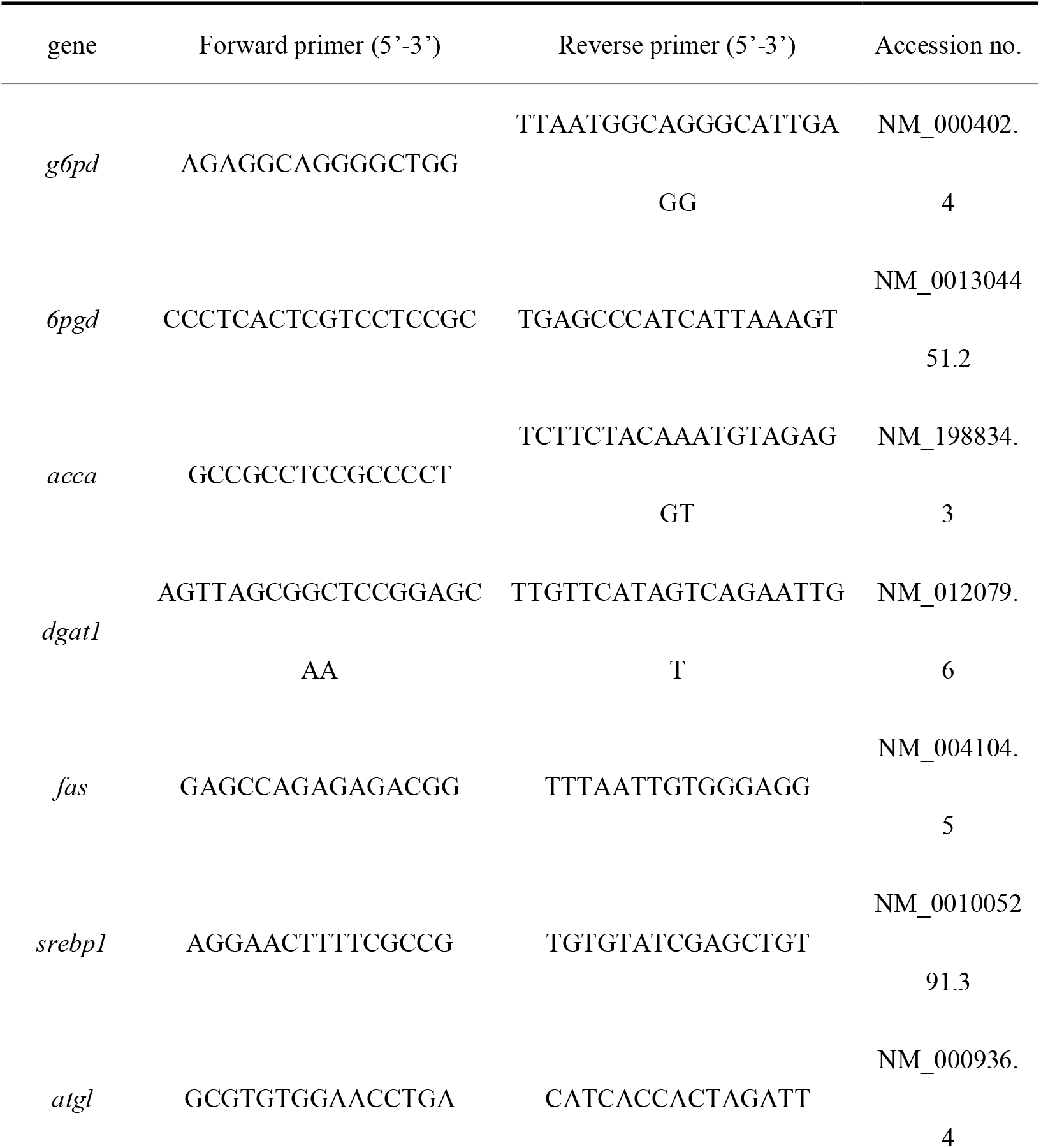
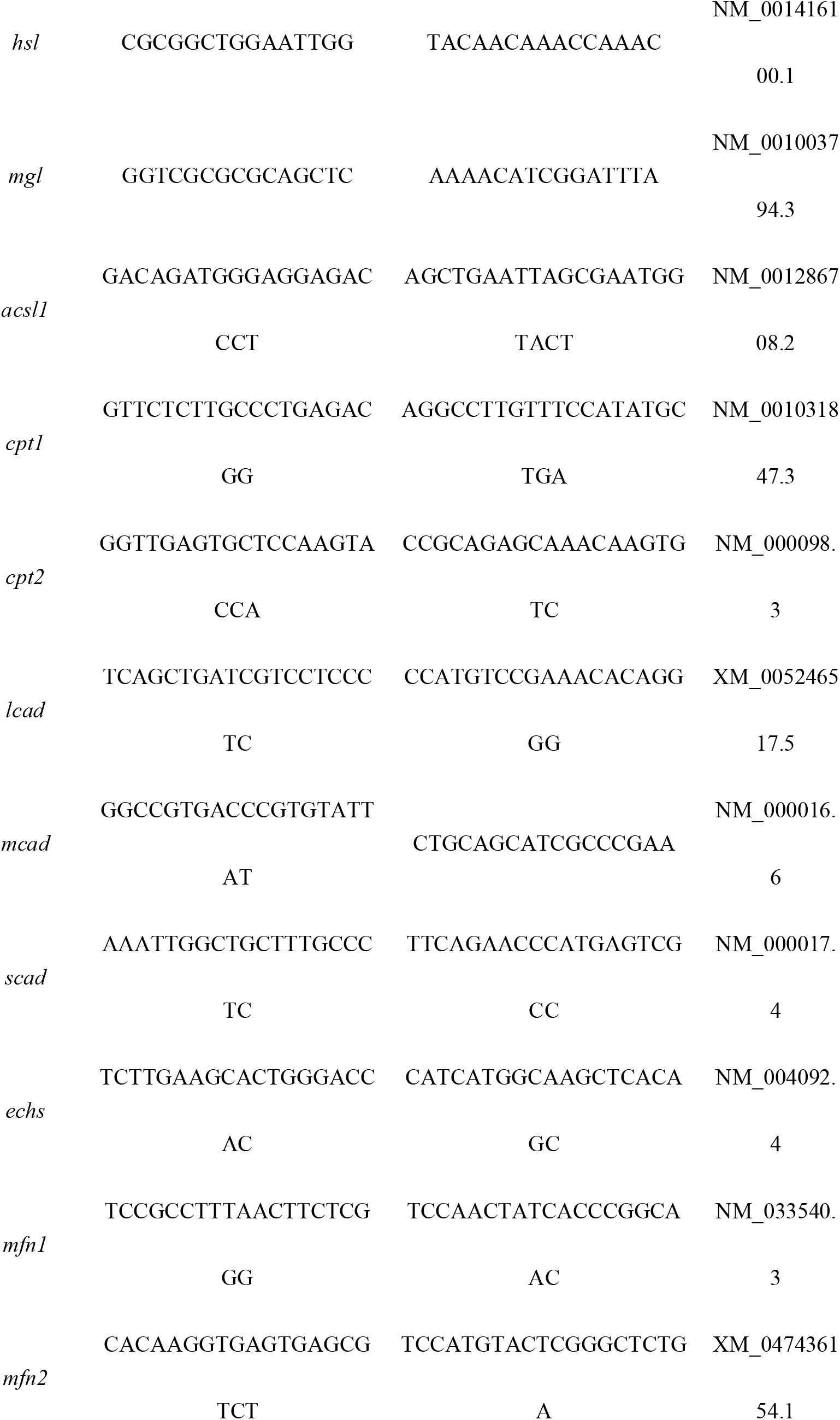
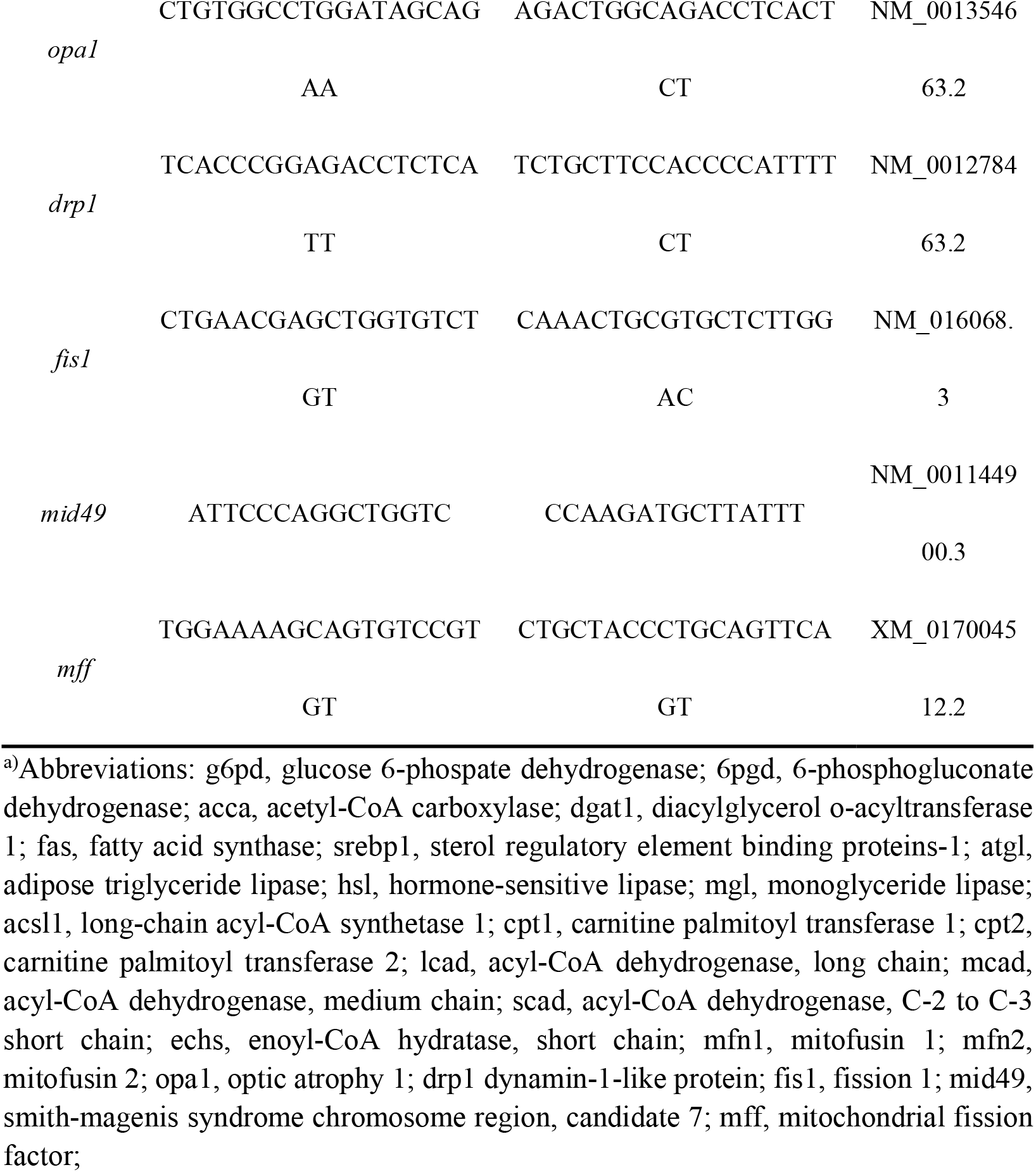
Primers in human for quantitative real-time PCR analysis.

**Table S8.**
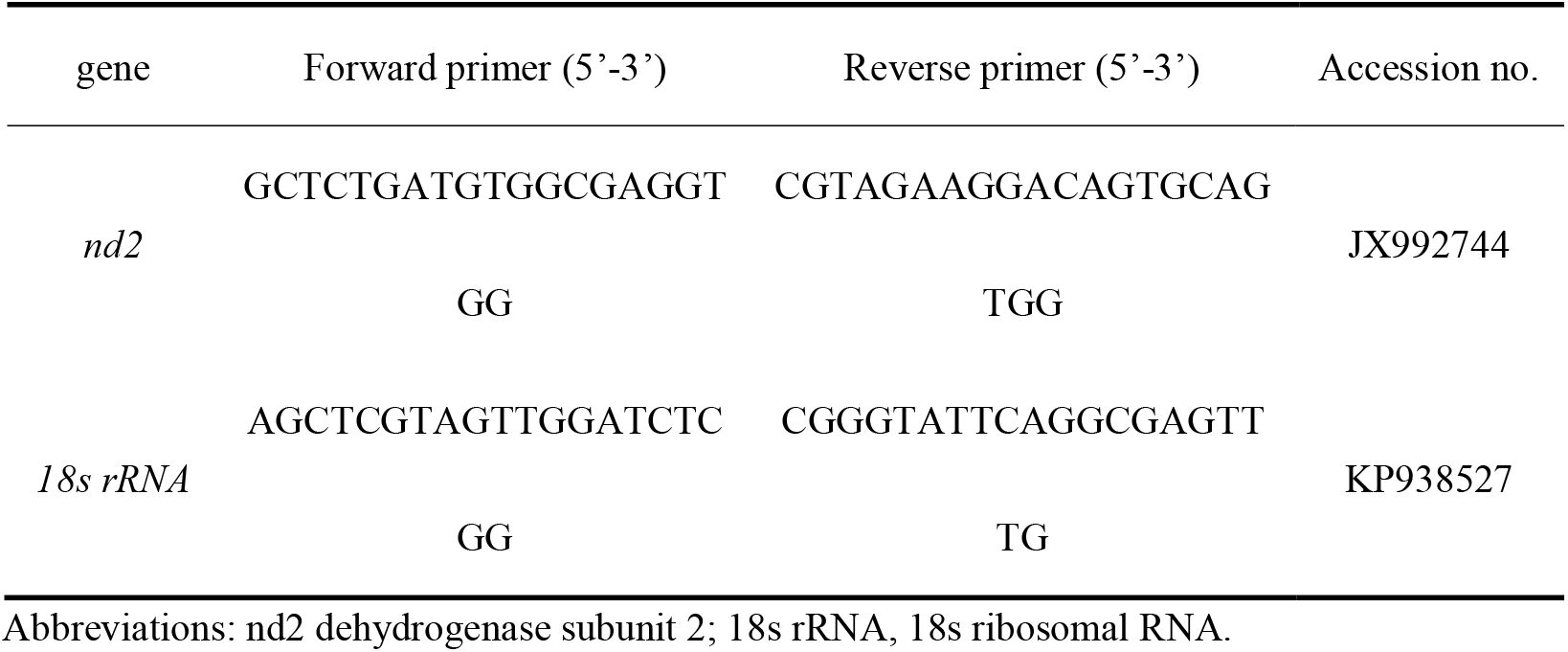
Primers for mtDNA copy number analysis.

**Table S9.**
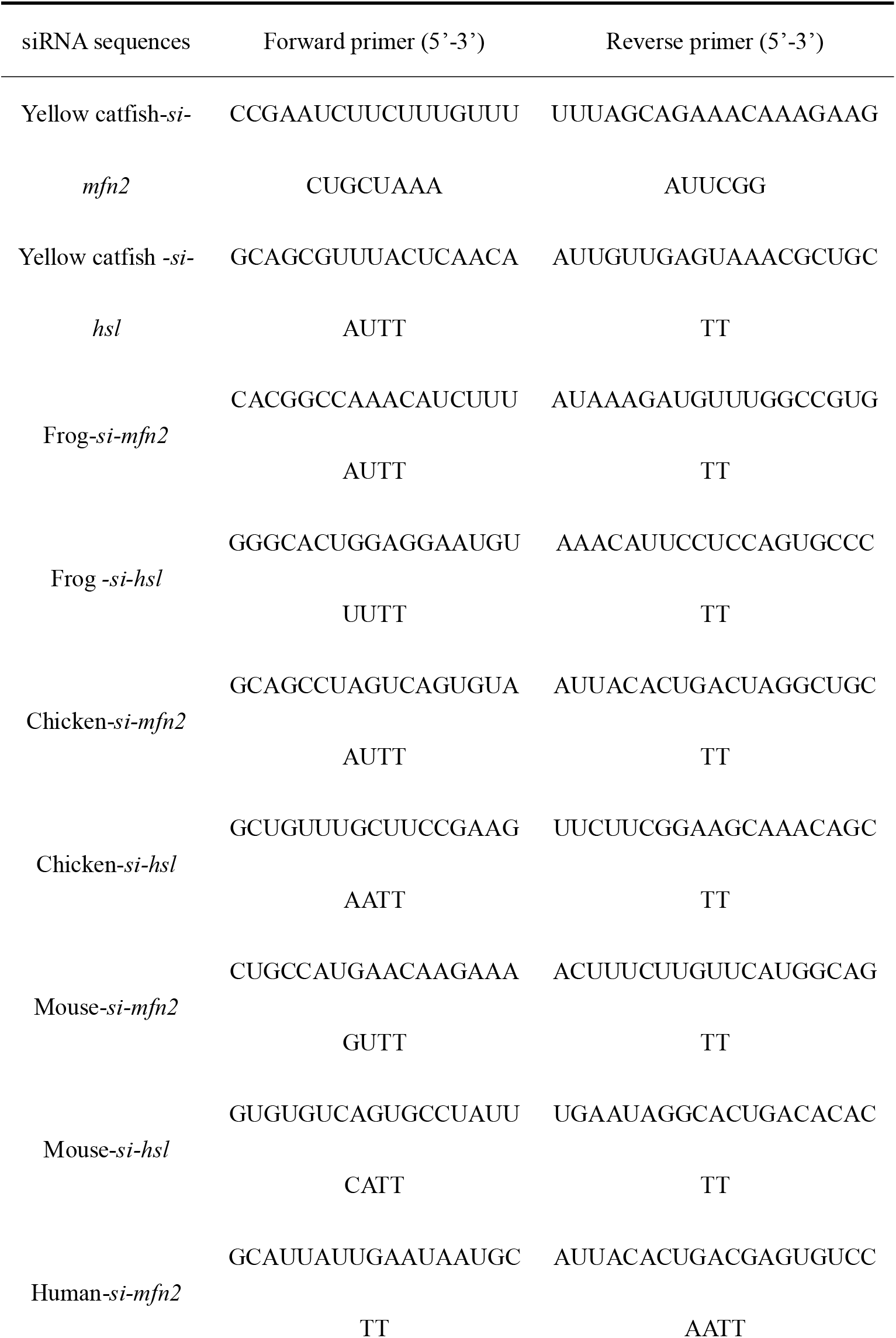
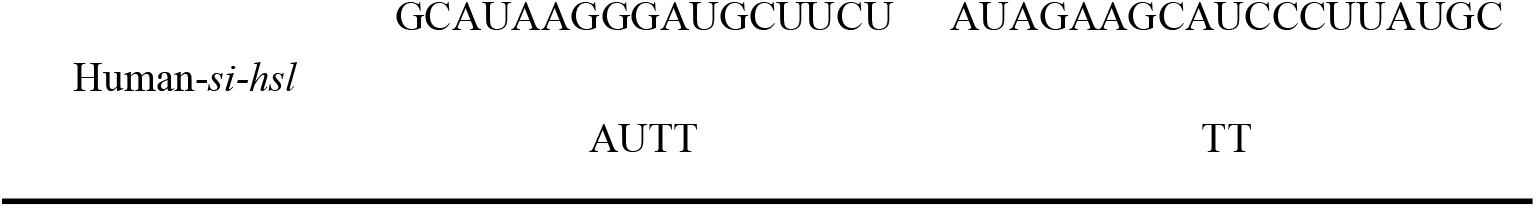
Primers used for siRNA sequences.

**Table S10.**
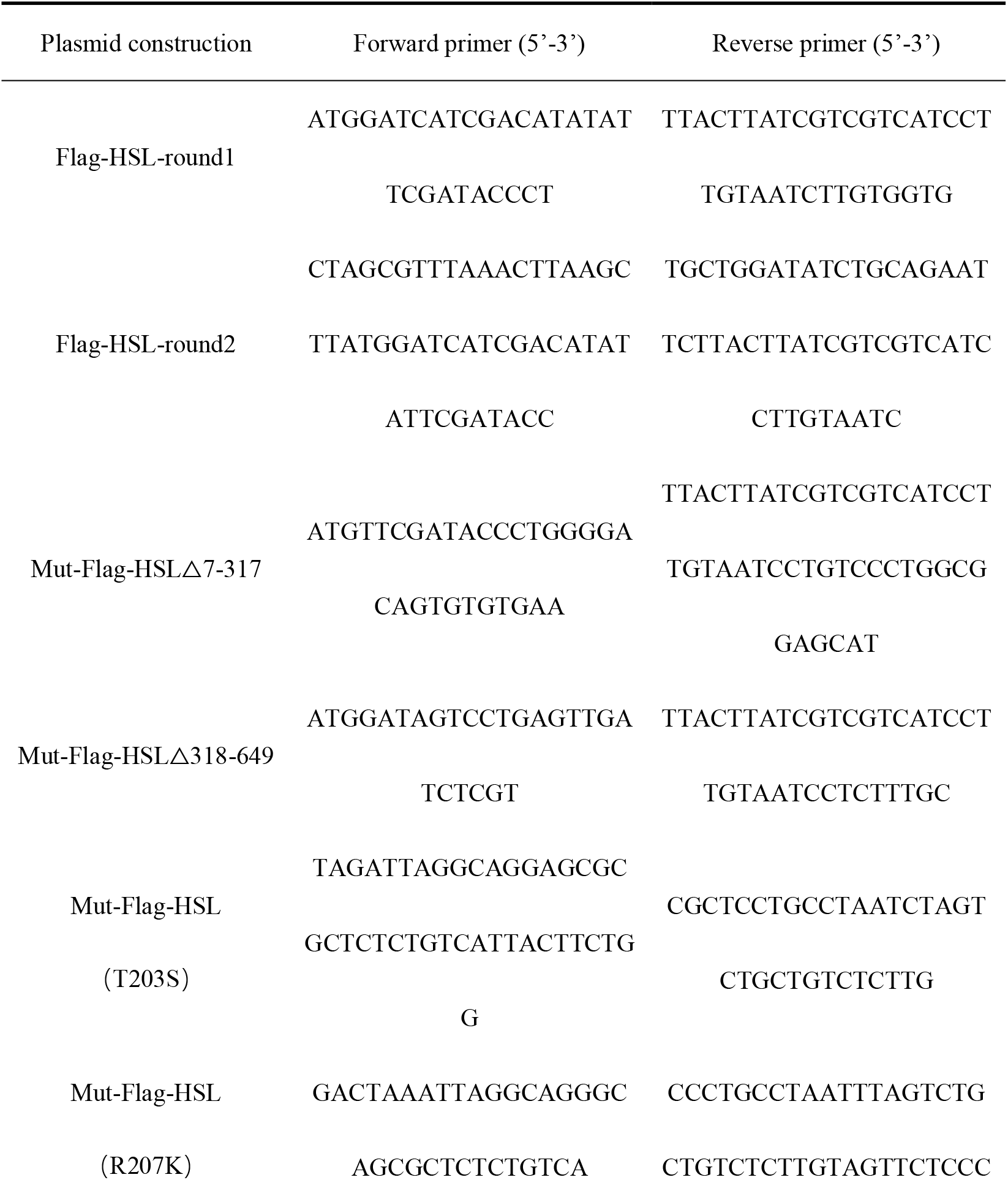
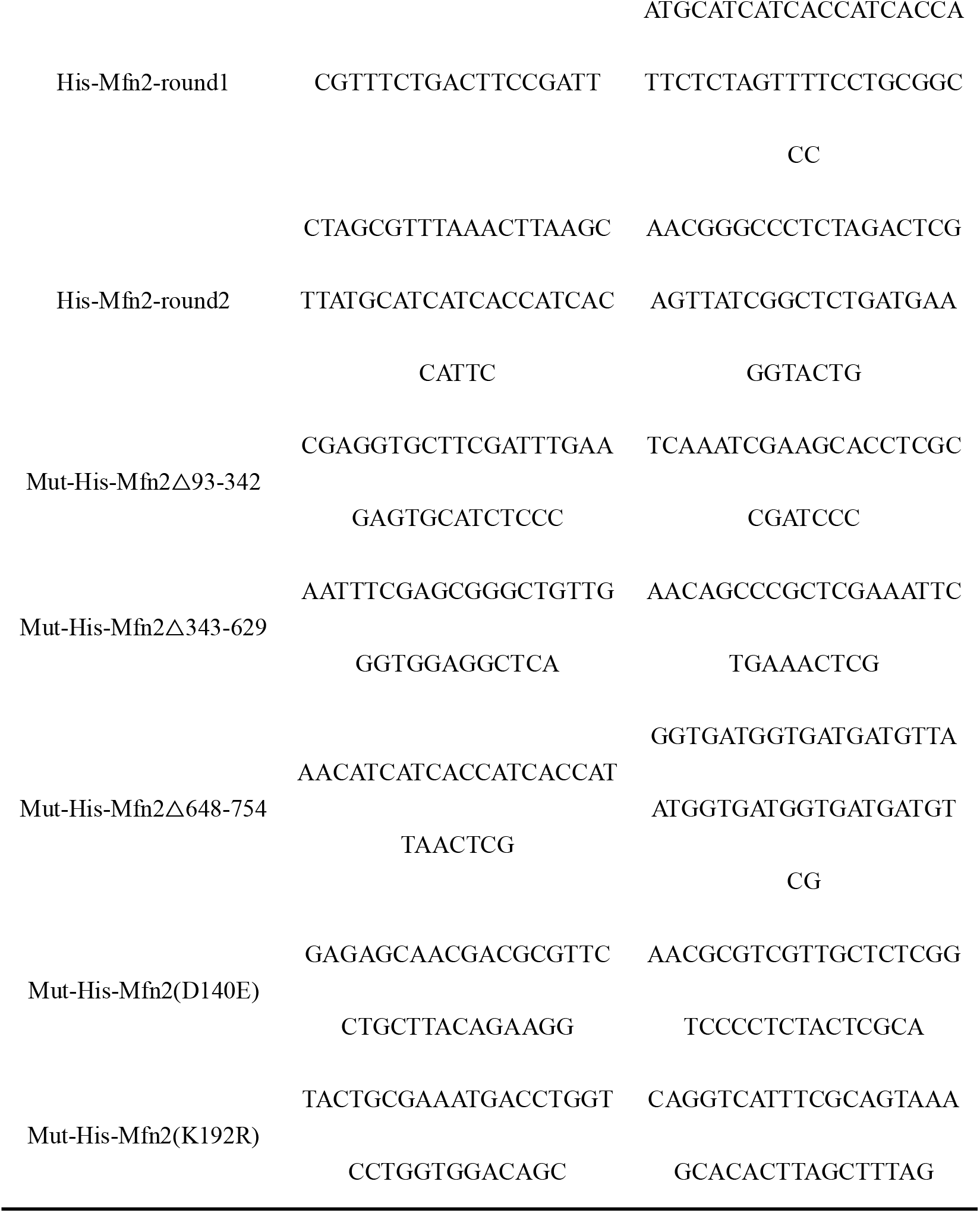
Primers used for plasmids construction.

**Table S11.**
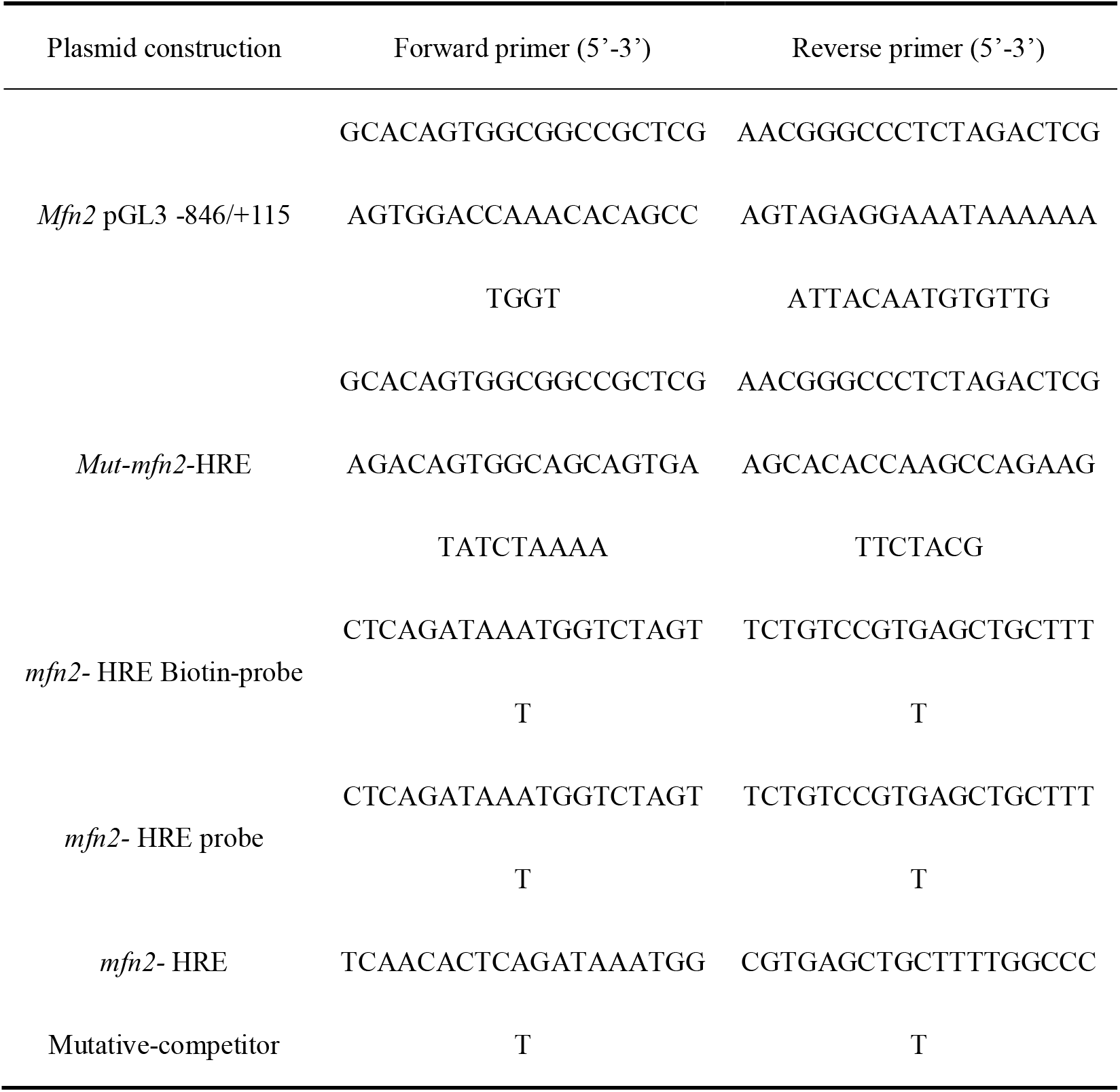
Primers used for mfn2 promoter cloning and Site-Directed Mutagenesis.

